# Mechanisms Matter: Transportability of Cellular Perturbation Effects

**DOI:** 10.64898/2026.05.08.723625

**Authors:** Shi-ang Qi, Paidamoyo Chapfuwa

## Abstract

Predicting cellular responses to perturbations across biological contexts is central to drug development, yet deep learning models often fail to outperform simple baselines. Leveraging causal transportability theory, we show that cross-context generalization is governed by shared causal mechanisms, not merely distributional similarity. To enable controlled evaluation, we develop a causal simulator that generates realistic semi-synthetic Perturb-seq datasets with tunable mechanistic divergence, providing benchmarks with known ground-truth causal structure. Further, we adapt the Vendi score to the perturbation setting to quantify effective response diversity and diagnose mode collapse, a failure mode invisible to standard per-perturbation metrics. Extensive experiments across six deep learning models and five simple baselines on semi-synthetic and real Perturb-seq datasets reveal a cross-context generalization gap: performance under cross-context splits drops substantially, often to simple baseline levels. Notably, even on synthetic data with fully specified causal structure, no model generalizes across contexts with different causal mechanisms. Across real and semi-synthetic datasets, lower cross-context concordance of transcriptome-wide impact tracks larger transfer gaps, providing an empirical proxy for perturbation-relevant mechanistic divergence. Code is available at https://github.com/microsoft/mechanisms_matter.

## 1 Introduction

A central goal of computational drug discovery is to predict how cells respond to genetic or chemical perturbations across diverse biological contexts, including cell types, tissues, and genetic backgrounds [Dixit et al., 2016, Hetzel et al., 2022]. Accurate cross-context perturbation response prediction could transform target identification and drug development by replacing costly exhaustive experimentation with principled in-silico prioritization [Bock et al., 2022, Roohani et al., 2025].

Large-scale perturbation atlases generated by Perturb-seq [Norman et al., 2019, Replogle et al., 2022, Zhu et al., 2025, Nadig et al., 2025] and related technologies now catalog single-cell transcriptomic responses to hundreds of perturbations. Yet the combinatorial space of perturbations (∼10^60^ chemical compounds, ∼20,000 genetic targets), cell types, donors, and disease states remains vastly under-sampled, limiting model generalization. Filling this space experimentally is infeasible: perturbation responses depend on cell type, differentiation stage, culture conditions, and genetic background, so measurements in one context do not automatically transfer to another. The Virtual Cell initiative [Roohani et al., 2025] seeks to bridge this gap through models trained on pooled multi-context data, but under what conditions such transfer is theoretically justified remains unclear.

A growing body of evidence indicates that increasing dataset size and model capacity has not translated into meaningful gains over simple baselines for perturbation response prediction [Bendidi et al., 2024, Ahlmann-Eltze et al., 2025, Csendes et al., 2025, Viñas Torné et al., 2025, Wenteler et al., 2025]. Two complementary explanations have emerged. First, standard per-perturbation metrics can be misleading. Whole-transcriptome Pearson correlation, for example, rewards alignment with the population-mean effect [Viñas Torné et al., 2025, Csendes et al., 2025]. More generally, scoring perturbations independently can miss mode collapse, in which models predict nearly identical responses across perturbations [Wu et al., 2025, Mejia et al., 2025]. The perturbation discrimination score (PDS) [Wu et al., 2025] detects this failure mode, but its *l*_1_ and *l*_2_ variants are scale-sensitive [Liu et al., 2025]. Second, models may lack the inductive biases needed to capture the causal mechanisms governing perturbation effects [Lorch et al., 2026, Dibaeinia et al., 2026, Wenteler et al., 2025]. Distinguishing among metric distortion, mode collapse, and model mis-specification is a prerequisite for meaningful progress.

Even when evaluation is corrected and models are made more expressive, formal guarantees of cross-context generalization remain absent [Dibaeinia et al., 2026]. Scaling data and architecture [Theodoris et al., 2023, Cui et al., 2024, Hao et al., 2024], disentangling perturbation effects [Lotfollahi et al., 2023, Piran et al., 2024, Zhang et al., 2024], matching distributions [Lotfollahi et al., 2019, Bunne et al., 2023, Dong et al., 2023, Adduri et al., 2026, Lorch et al., 2026, Klein et al., 2025, Palla et al., 2026], and incorporating biological priors [Roohani et al., 2024, He et al., 2025] have each shown promise within individual contexts. Yet the cross-context generalization gap itself has not been systematically quantified. Causal transportability [Bareinboim and Pearl, 2013] offers a principled framework for determining whether and how perturbation effects generalize across contexts, including when mechanism shifts are identifiable and adjustable through a valid transport formula. Because causal cascades mediating perturbation effects are rarely fully characterized in biology [Amit et al., 2009], we treat mechanism invariance as a sufficient but not necessary guiding principle, emphasizing that distributional similarity alone does not guarantee transportability (Figure 1).

**Figure 1:**
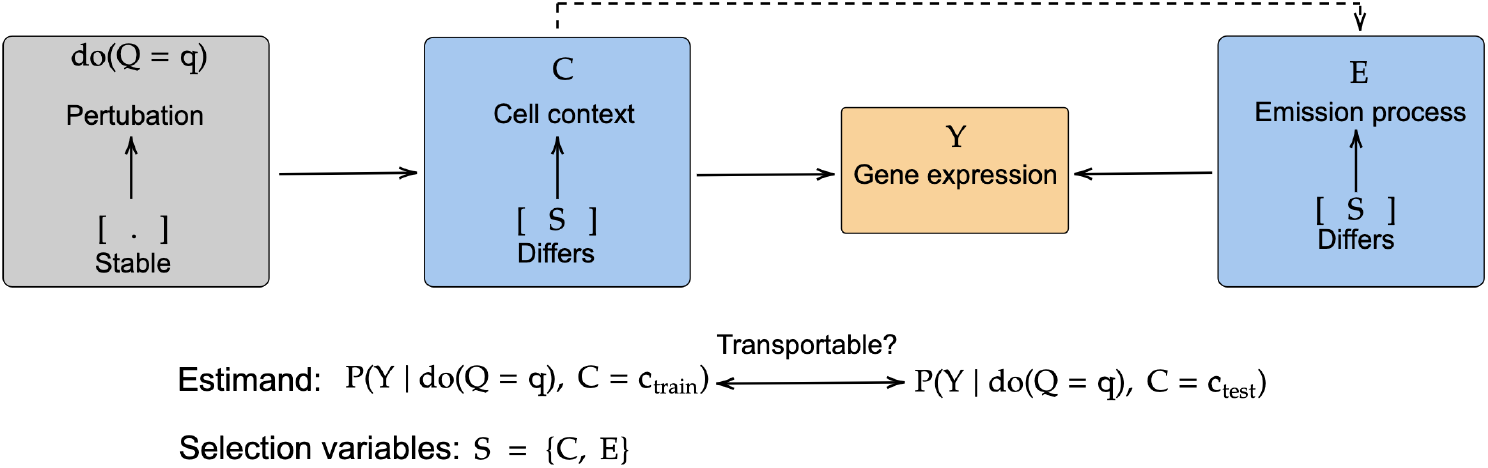
Overview of the transportability problem in cellular perturbation prediction. Given training pairs {(*q, c*_train_), (*q*^*′*^, *c*_test_) : *q*^*′*^ ≠ *q*}, the goal is to predict *P* (*Y* |do(*Q* = *q*), *C* = *c*_test_) for the *unseen (perturbation, context) pair* (*q, c*_test_), where *Y* denotes the cellular response and do(*Q* = *q*) assigns perturbation *q* [Pearl, 2009]. *The selection variables S* = {*C, E*} denote biological context and emission process, respectively. Observed components of *E* (*e*.*g*., library size and batch) can be adjusted for, whereas latent components (*e*.*g*., dropout rate and technical noise) induce domain shift.

To address these challenges, we make the following contributions:

- We formally connect perturbation prediction to causal transportability [Bareinboim and Pearl, 2013], showing that cross-context generalization is governed by shared causal mechanisms, not distributional similarity.
- We extend the framework of Lorch et al. [2026] into a cross-context causal simulator with realistic semi-synthetic data, known causal structure, and tunable mechanistic divergence, enabling controlled evaluation of generalization.
- We adapt the Vendi score [Friedman and Dieng, 2023] as a diagnostic of effective response diversity. Together, the Vendi score and PDS [Wu et al., 2025] characterize perturbation-response geometry in terms of effective diversity and discrimination.
- Across six deep learning models and five simple baselines on semi-synthetic and real Perturb-seq datasets, we reveal a cross-context generalization gap that widens as transcriptome-wide impact (TI; [Nadig et al., 2025]) concordance decreases, supporting TI concordance as an empirical proxy for mechanistic divergence across contexts.

## 2 Perturbation transportability as a causal problem

We formally frame the virtual cell challenge [Roohani et al., 2025] as a causal transportability problem: under what conditions can cellular perturbation effects observed in one biological context be validly transferred to another? Leveraging causal transportability theory [Bareinboim and Pearl, 2013] and a semi-synthetic data-generating process, we derive necessary and sufficient conditions for such transfer within the linear regime and show that when specific *causal mechanisms* differ across contexts, unadjusted transfer is biased, and recovery is only possible if a context-specific transport map is identifiable from the available data.

### 2.1 Problem formulation

Let *Y* ∈ ℤ^*G*^ denote gene-expression counts for *G* genes, *Q* ∈ {0, 1, …, *P*} the perturbation index (*Q*=0 for control), and *C* the biological context (*e*.*g*., cell type, tissue, or experimental condition). Perturbation prediction aims to infer

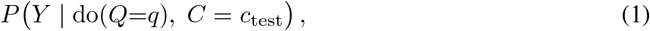

where do(*Q*=*q*) denotes an intervention that assigns perturbation *q* [Pearl, 2009]. *We estimate this* distribution for an *unseen (perturbation, context) combination*, using interventional samples from the observed combinations {(*q, c*_train_)}∪ {(*q*^*′*^, *c*_test_) : *q*^*′*^ ≠ *q*} and target-context control samples drawn from *P* (*Y* do(*Q*=0), *C*=*c*_test_). For pseudobulk prediction, the estimand reduces to the interventional mean E [*Y* | do(*Q*=*q*), *C* = *c*_test_]. We illustrate the assumed selection diagram for cellular perturbation transportability in Figure 1. The emission process *E* governs the mapping from the latent state to observed scRNA-seq counts and may itself vary across context, yielding selection variables *S* = {*C, E*} . We assume two key unobserved mechanisms that may differ across cellular contexts:

i. the causal influence matrix *A*_*C*_ ∈ ℝ^*G×G*^, a linear approximation of the active gene regulatory network (GRN) in context *C*: entry [*A*_*C*_]_*ij*_ ≠ 0 indicates that gene *j* directly regulates gene *i* (positive: activation, negative: repression);
ii. the basal expression vector *B*_*C*_ ∈ ℝ^*G*^, encoding the context-specific baseline expression program, reflecting the combined influence of transcription rates, epigenetic state, chromatin accessibility, and lineage-specific regulatory activity [Huang et al., 2005, Klemm et al., 2019].

A context shift alters *A*_*C*_, *B*_*C*_, or both: *A*_*C*_ rewires the regulatory program and thus the perturbation response (cell-type and lineage differences), while *B*_*C*_ shifts the basal state (batch and activation-state variation), leaving the latent response mechanism intact but potentially altering measured effects under nonlinear emission. Transportability depends on *which* of these differ across contexts and whether the differences are identifiable from the available data. In the trivial case *c*_train_ = *c*_test_, the problem reduces to interpolation over seen (perturbations, context) combinations, and transportability holds by construction. Throughout, let 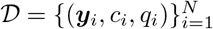 denote a dataset of *N* observations.

### 2.2 Causal simulator with tunable transportability

We generate semi-synthetic scRNA-seq data from a continuous-time linear stochastic differential equation (SDE) in a latent gene-expression space, followed by a negative binomial observation model. We term this simulator CausalDGP, see Figure 2, for an illustration of the assumed selection diagram. Following Lorch et al. [2026], we adopt the Ornstein–Uhlenbeck (OU) process [Uhlenbeck and Ornstein, 1930], which is *causally structured, stationary, and analytically tractable*: these properties allow us to derive closed-form transportability conditions while generating realistic gene expression counts through a negative binomial likelihood.

**Figure 2:**
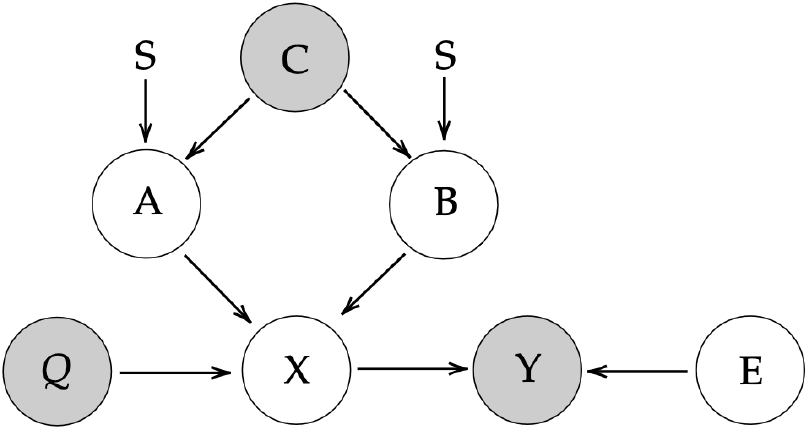
Selection diagram of the assumed CAUSALDGP, with unobserved mechanistic, context-specific selection variables *S* = {*A, B*} . Cell context *C*, perturbation *Q*, and gene expression counts *Y* are observed. The unobserved emission process *E*, mapping the latent state *X* to observed counts *Y*, is invariant across contexts.

**Figure 3:**
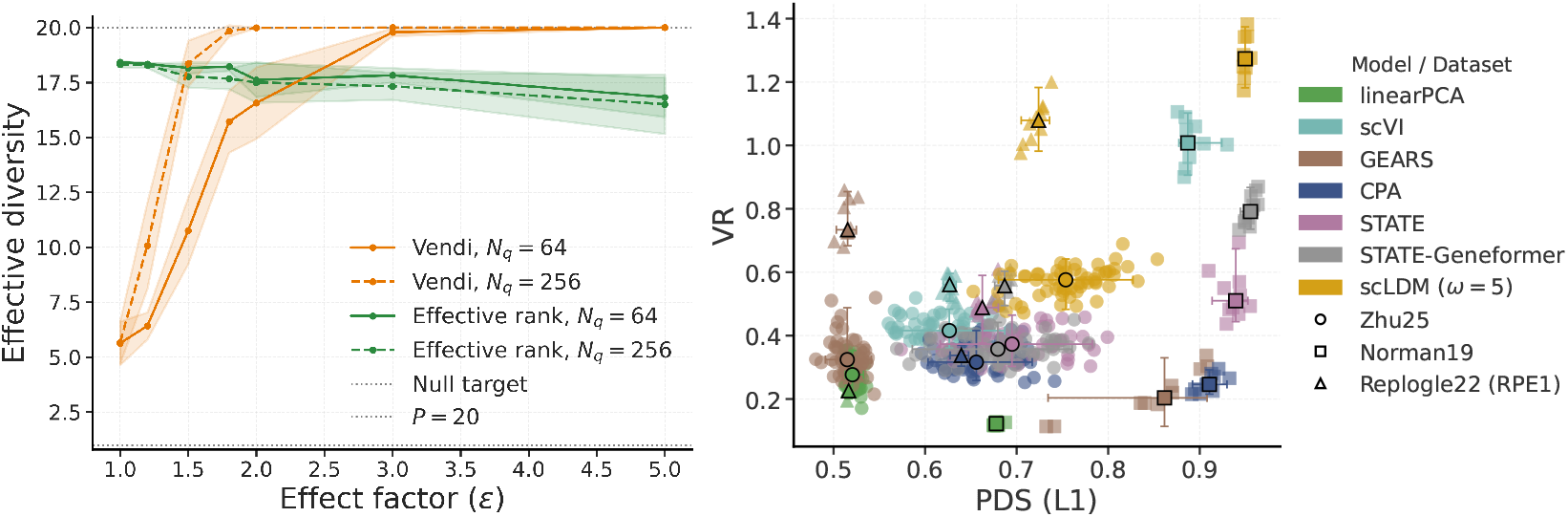
(a) Effective diversity in synthetic DIRECTDGP datasets (*P* = 20; *p*_effect_ = 0.1; *β* = 0). Lines show means over five replicates (shading: *±* 1 SD); solid/dashed lines indicate *N*_*q*_ = 64*/*256. (b) VR vs. PDS (*l*_1_) across models and datasets. Color and shape indicate model and dataset; enlarged markers and error bars show medians and 95% percentile intervals per model–dataset pair.

**Figure 4:**
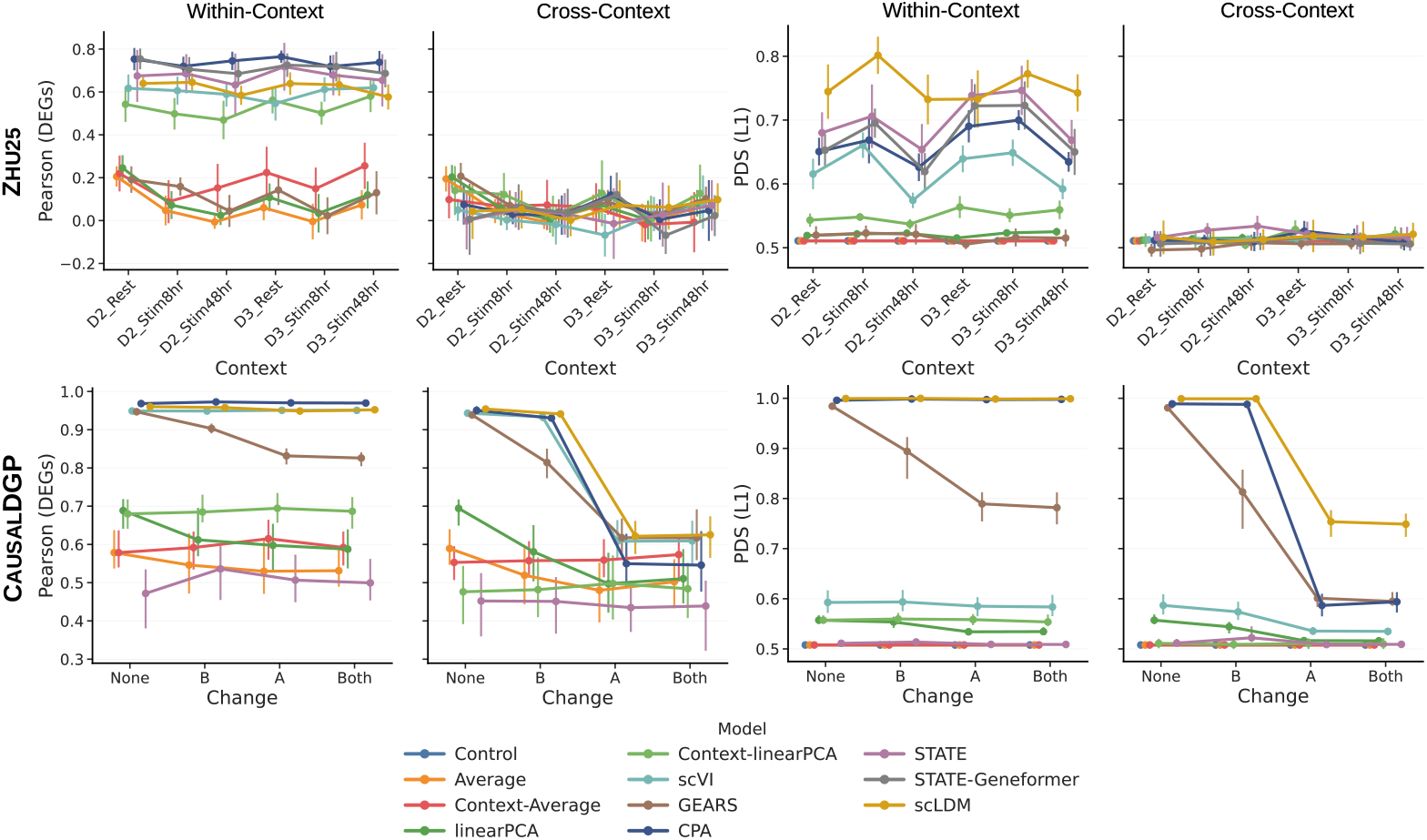
Cross-context gaps on Zhu25 (top) and CausalDGP (bottom) for DEG-restricted Pearson ***δ*** and PDS (*l*_1_). The *x*-axis shows six Zhu25 test contexts (2 donors *×* 3 activation states) and four CAUSALDGP mechanism modes (None, A, B, Both). We plot medians and interquartile ranges. scLDM is evaluated with classifier-free guidance weight *ω* = 5.

Specifically, we model the latent expression state *X*(*t*) ∈ ℝ^*G*^ of a cell with *G* genes as a multivariate OU process [Lorch et al., 2026]:

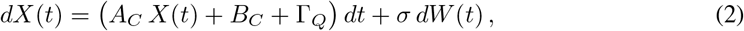

where *A*_*C*_ ∈ ℝ^*G×G*^ is a sparse causal influence matrix encoding the GRN under biological context *C, B*_*C*_ ∈ ℝ^*G*^ is a context-specific baseline drift bias, *Q* ∈ {0, 1, …, *P*} indexes the perturbation condition, and Γ_*Q*_ ∈ ℝ^*G*^ is a sparse shift vector encoding the direct effect of perturbation *Q* (with Γ_*Q*_ = **0** for the unperturbed control *Q* = 0), *dW* (*t*) denotes *G*-dimensional Brownian motion, and *σ* > 0 is a diffusion coefficient encoding *intrinsic biological stochasticity*. We refer to Lorch et al. [2026] for detailed construction of *A*_*C*_. Provided *A*_*C*_ is Hurwitz (all eigenvalues with strictly negative real part), the process admits a unique stationary Gaussian distribution

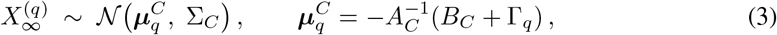

where the covariance Σ_*C*_ is the unique positive-definite solution of the continuous Lyapunov equation 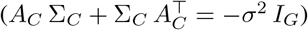.

#### Emission process

In practice, the latent states *X*(*t*) are *unobserved*. Given a stationary sample 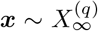, observed counts are generated independently per gene through an emission process *E*, modeled as a negative binomial (NB) likelihood, capturing both *technical measurement noise* and *intrinsic transcriptional variability*:

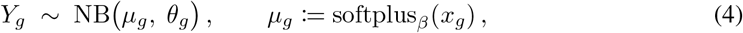

where *µ*_*g*_ is the mean parameter and *θ*_*g*_ is a gene-specific inverse-dispersion. The softplus link softplus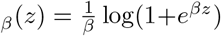 maps real-valued latent states to non-negative intensities. The negative binomial captures intrinsic transcriptional noise arising from bursty gene expression [Amrhein et al., 2019], and has become the standard observation model for scRNA-seq [Lopez et al., 2018].

Collectively, the emission process *E* is fully specified by the gene-wise dispersion parameters 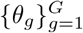 and the softplus link, which defines the mapping from latent state to observed counts. As illustrated in the selection diagram (Figure 2), we assume *E* is invariant across contexts (*E*_*C*_ = *E* for all *C*), so all non-transportability in the semi-synthetic data arises from the latent dynamics alone. In practice, context shifts in library size, capture efficiency, or dispersion would introduce additional measurement-level non-transportability not modeled here. Following Mejia et al. [2025], we set 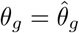 where 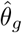 is estimated from the Norman et al. [2019] dataset via maximum likelihood, ensuring realistic count statistics while retaining full control over the latent causal structure. Complete details are provided in Appendix B.2.

### 2.3 Transportability conditions

Consider two biological contexts (*e*.*g*., cell types) *C* and *C*^*′*^ with potentially distinct causal influence matrices *A*_*C*_, *A*_*C*_*′* and baseline biases *B*_*C*_, *B*_*C*_*′*, but a shared perturbation Γ_*q*_. From (3), the perturbation effect in each context is

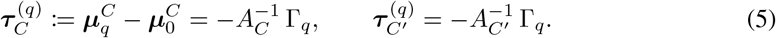

#### Proposition 1

**(Direct transportability of perturbation effects)**

*From (5) the effect of perturbation q is directly transportable from C to C*^*′*^ *without context-specific adjustment if and only if*

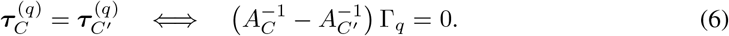

Crucially, this condition is *perturbation-specific*: for perturbation *q*, it involves only the columns of 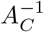 and 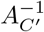 indexed by the support of Γ_q_. Thus, a pair of contexts may permit direct latent transport for one perturbation but not another. Marginal divergence between *P*_*C*_(*Y* ) and *P*_*C*_*′* (*Y* ) neither establishes nor precludes direct latent transport: even when 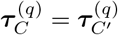, nonlinear emission can yield different measured perturbation effects across baselines *B*_*C*_ and *B*_*C*_*′*, while differences in *σ* and irrelevant pathways in *A* may further shift observed distributions. Thus, perturbation-relevant mechanism invariance, unlike overall distributional similarity, is sufficient for direct latent transport.

More generally, effect equality is unnecessary if a context-specific transport map is identifiable. When Γ_q_ is shared across contexts, then 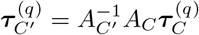. Thus, unequal effects are transportable when 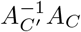 is identifiable; otherwise, the held-out target effect cannot generally be recovered. Typical Perturb-seq screens (*P* ≪ *G*) can identify 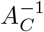 only on span {Γ_*q*_} ; full recovery requires *G* independent perturbations. Even then, noise, variable perturbation efficiency, and nonlinear dynamics make single-cell GRN inference challenging and approximate [Pratapa et al., 2020].

## 3 Metrics

### Algorithm 1

Null-Calibrated Pseudobulk Vendi Score

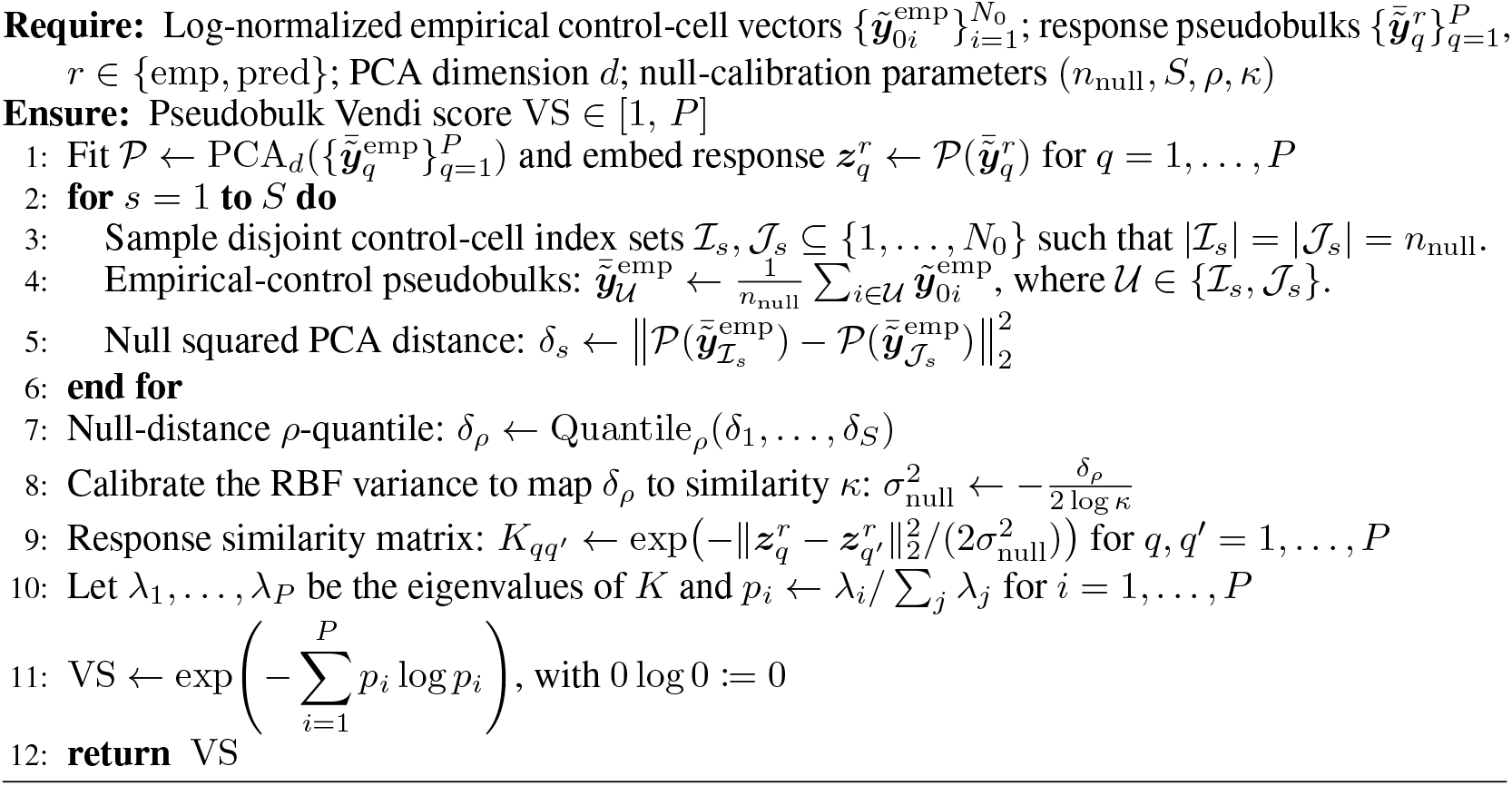

In practice, most models do not infer the causal matrix *A*_*C*_ in (5), and the ground-truth perturbation effect is typically unknown. Deep learning models approximate the full conditional distribution *Pϕ* (Y | do(*Q*=*q*),*C*)?, from which predicted means are obtained, while simple baselines directly estimate E [*Y* | do(*Q*=*q*), *C*] by averaging observed training samples. Our focus is on quantifying the *cross-context generalization gap*: the degradation in perturbation-effect prediction when models transfer from observed to unseen biological contexts. We therefore use perturbation-effect metrics that capture complementary aspects of this gap. In both cases, we define the predicted perturbation effect as

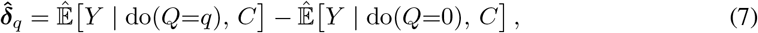

where 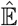 denotes the model’s predicted mean expression, *a*.*k*.*a* pseudobulk, whether derived from a learned distribution or from empirical averaging. This formulation requires only predicted means for both the perturbed and control conditions, making downstream metrics applicable to all models. Below, we describe the proposed Vendi score for quantifying perturbation-response diversity. Additional metrics are detailed in the Appendix, spanning three families: *reconstruction metrics* including distributional (MMD, FD, JSD) and point-wise (MSE, MAE) measures (Appendix D.4); *perturbation effect metrics* including Pearson ***δ*** (14), the perturbation effect discrimination score (PDS) (13), coefficient of determination 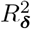 (15), and transcriptome-wide impact (TI) (19) (Appendix D.2); and *differentially expressed gene (DEG) set metrics* including Jaccard index, recall, and precision (Appendix D.3). Because per-perturbation metrics are sensitive to control bias and perturbation-gene sparsity (Appendix E.6), we report DEG-restricted variants throughout.

### 3.1 Effective perturbation response diversity: Vendi score

Per-perturbation metrics such as Pearson ***δ*** evaluate predictions independently, potentially masking mode collapse [Wu et al., 2025] or reflecting signal strength rather than model quality for sparse effects (Appendix E.6). We therefore quantify effective response diversity using a *null-calibrated* pseudobulk Vendi score adapted from Friedman and Dieng [2023], which converts pairwise distances in an empirical-reference PCA space into RBF similarities (Algorithm 1).

We calibrate the RBF scale using pairs of disjoint pseudobulks sampled from empirical control cells, each containing *n*_null_ cells. We choose the scale such that the *ρ*-quantile of their squared PCA distances corresponds to a target similarity *κ* (Table 8, Appendix). This calibration captures finite-sample variability at the reference size *n*_null_ but does not explicitly adjust for perturbation-specific sample sizes *N*_*q*_. We limit extreme mismatch by requiring *N*_*q*_ *≥* 64. The Vendi score is the exponential spectral entropy of the resulting perturbation-similarity matrix and represents the effective number of distinct perturbation responses. We compute empirical and predicted scores using the same empirical PCA transformation and null calibration and report the *Vendi Ratio*:

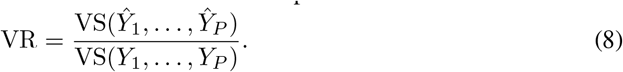

Here, VS denotes the Vendi score, and *Y*_*q*_ and *Ŷ*_*q*_ denote the empirical and predicted perturbation responses, respectively. The raw predicted Vendi score is unnormalized, whereas the Vendi Ratio normalizes predicted diversity by empirical diversity. A ratio near one indicates preserved empirical diversity, although finite-sample noise may cause an optimal deterministic predictor to yield VR *<* 1. Complete collapse of the predicted responses gives VS(*Ŷ*) = 1 and hence VR = 1*/*VS(*Y* ), which equals 1*/P* only when the empirical responses attain maximal diversity.

To characterize complementary aspects of *perturbation-response geometry*, we report the Vendi score alongside PDS. PDS measures response distinguishability, whereas the Vendi score quantifies effective diversity by accounting for redundancy. Together, they assess whether responses are both distinct and nonredundant.

## 4 Experiments

All code is implemented in Python and is publicly available at https://github.com/microsoft/mechanisms_matter.

### 4.1 Baselines

We evaluate six deep learning models spanning *transport-based* (STATE [Adduri et al., 2026], scLDM [Palla et al., 2026]), *graph-based* (GEARS [Roohani et al., 2024]), *disentanglement-based* (CPA [Lotfollahi et al., 2023], scVI [Lopez et al., 2018]), and *foundation-model* approaches (STATE-Geneformer, combining Geneformer [Theodoris et al., 2023] embeddings with a STATE decoder). We also evaluate five simple baselines: a global control mean, plus both global and context-specific variants of perturbation mean [Csendes et al., 2025, Kernfeld et al., 2023] and linearPCA [Ahlmann-Eltze et al., 2025]. We adapt scVI to perturbation prediction using our transport formula. To predict *unseen (perturbation, context) combinations*, we use STATE’s raw-count state-transition variant rather than its pretrained embedding model. Geneformer’s pretraining predates Zhu25 and excludes immortalized cell lines, making our benchmark profiles unseen despite including primary immune cells. GEARS uses gene ontology and training-derived co-expression graphs; for synthetic data, which lack an external ontology, we use only the latter. See Tables 3 and 4 in Appendix A for a summary of the considered baselines and their detailed transport formulae, respectively.

### 4.2 Datasets

We focus on Perturb-seq because its targeted gene interventions provide an experimental analog of Pearl’s do(*·*) operator [Pearl, 2009] for studying causal gene function at single-cell resolution. Extending the framework to cytokines and chemical perturbations [Klein et al., 2025] remains an important direction for future work. See Table 1 for summary statistics and Appendix C.1 for complete details.

**Table 1:** Summary of the cellular perturbation datasets after preprocessing. HVGs: Highly variable genes.

| Statistic | ZHU25 | REPLOGLE22/NADIG25 | NORMAN19 | CAUSALDGP | DIRECTDGP |
| --- | --- | --- | --- | --- | --- |
| Cell type | Primary CD4+ T cells | RPE1-K562 /<br>HepG2-K562 /<br>Jurkat-K562 | K562 | - | - |
| Perturbation type | CRISPRi knockdown | CRISPRi knockdown | CRISPRa over-expression | synthetic | synthetic |
| Control cells $N_0$ | 393,216 | 22,176 / 15,667 / 22,704 | 11,855 | 1,024 | 128–1,024 |
| Perturbed cells $N_P$ | 203,870 | 266,087 / 144,160 / 300,954 | 71,742 | 1,024 × 128 | (128–512) × (20–50) |
| HVGs $G$ | 3,463 | 7,226 / 7,595 / 7,563 | 8,192 | 128 | 1,024–8,192 |
| Perturbations $P$ | 91 | 827 / 479 / 983 | 156 | 128 | 20–50 |
| Total contexts $C$ | 12 (4 donors × 3 conditions) | 2 / 2 / 2 | 1 | 2 | 1 |

#### Semi-synthetic benchmarks

We use two data generating approaches. DirectDGP, adopted from Mejia et al. [2025], simulates scRNA-seq perturbation data within a single context using a negative binomial model with parameters estimated from the Norman19 dataset. We use it to study metric behavior, including metric distortion and Vendi score validation, in controlled within-context settings. CausalDGP (Section 2.2) extends this to two contexts with distinct latent causal mechanisms (*A*_*C*_, *B*_*C*_). By varying which mechanisms differ across contexts (`None`, `A`, `B`, `Both`), it enables control over whether perturbation effects are transportable by construction. Refer to Appendix B for complete simulation details.

#### Validating

CausalDGP Figure 6 (Appendix) illustrates cell-level heterogeneity and context-dependent shifts in expression space. Beyond this qualitative visualization, we evaluate marginal fidelity, as emphasized in prior simulator studies [Zappia et al., 2017, Sun et al., 2021], and down-stream task behavior. We compare CausalDGP with real Perturb-seq data within each cellular context using marginal, co-expression, perturbation-effect, effective-diversity, and discriminability metrics (Table 6, Appendix). Given the heterogeneity of real Perturb-seq data, CausalDGP falls within observed ranges for dispersion (CoV), sparsity (Zero), co-expression (Frac|*r*|>0.3), and effect size 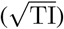. Across scenarios, Vendi reaches 20.1–27.2% of its theoretical maximum, *P* = 128, indicating substantial effective diversity. Given the simulator design, high split-half PDS is expected by construction and serves as a sanity check that sampling noise does not obscure the simulated perturbation effects.

**Figure 5:**
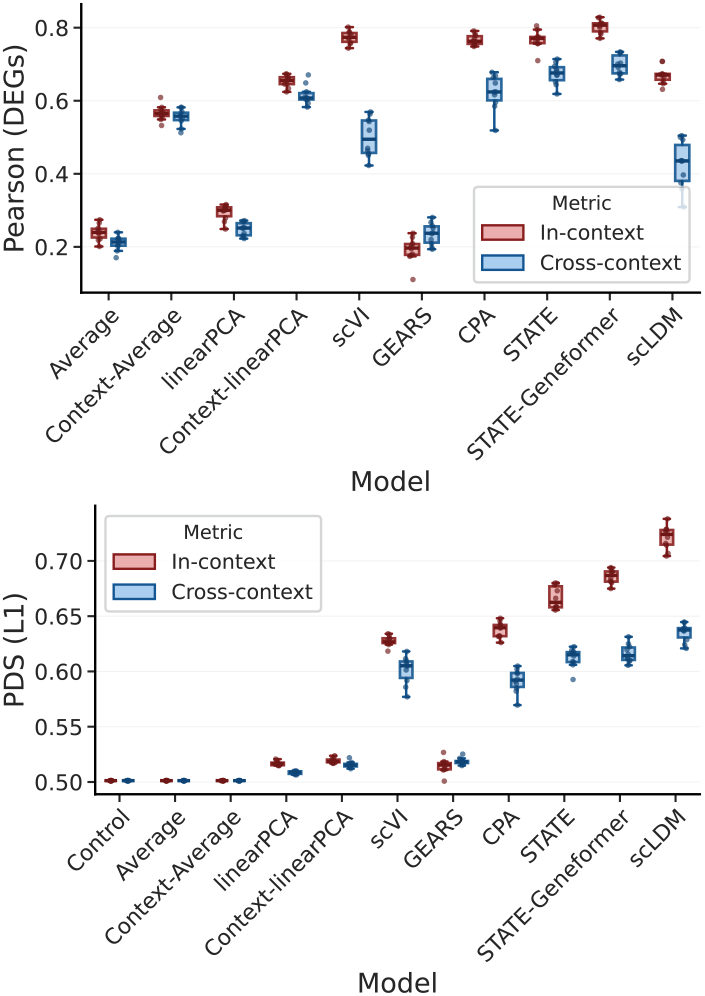
Cross-context generalization gap on Replogle22/Nadig25 for Pearson ***δ*** (DEG-restricted) and PDS (*l*_1_). Models are trained on RPE1 and evaluated on held-out K562 perturbations. scLDM is evaluated with classifier-free guidance weight *ω* = 5.

**Figure 6:**
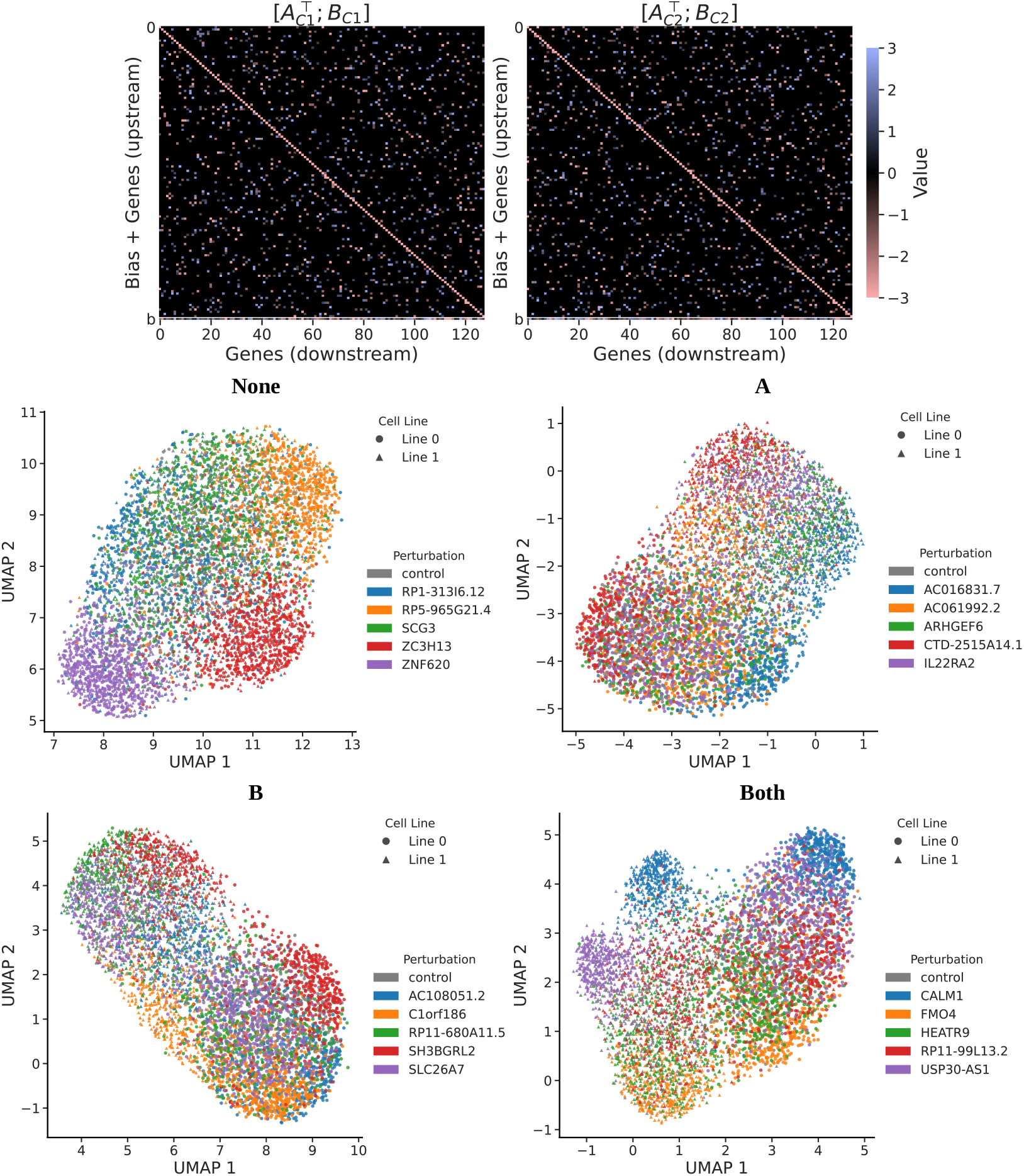
Qualitative visualization of CausalDGP under the four mechanism-diversity modes. **Top row:** context-specific latent mechanisms for the two simulated contexts, shown as concatenated parameter blocks 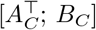, where *A*_*C*_ is the sparse signed interaction matrix and *B*_*C*_ is the baseline drift vector. **Bottom row:** UMAP projections of simulated cells under each diversity mode (`None`, `A`, `B`, `Both`), with 128 genes, 5 perturbations plus control, and 1,024 cells per condition. UMAP was computed on the top 50 principal components of the raw count matrix. Color denotes perturbation identity, and marker shape denotes context *C* ∈ {0, 1}.

#### Real-world datasets

We evaluate on three Perturb-seq datasets spanning progressively more challenging transportability settings:

- Norman19 [Norman et al., 2019]: single- and double-gene perturbations in K562 cells (one context), used to study metric distortion and Vendi-score sensitivity.
- Replogle22/Nadig25 [Replogle et al., 2022, Nadig et al., 2025]: single-gene perturbations across K562, RPE1, Jurkat, and HepG2 cell lines (four contexts). K562 and RPE1 are from Replogle et al. [2022], while Jurkat and HepG2 are from Nadig et al. [2025]. These lines contrast hematopoietic (K562, Jurkat) with epithelial (RPE1, HepG2) and cancer with non-transformed (RPE1), providing a test bed across genuine biological divergence. We treat the partner line as the source context and K562 as the target (evaluation), defining one transportability task per pair.
- Zhu25 [Zhu et al., 2025]: large-scale primary human CD4+ T-cell Perturb-seq across 12 contexts (4 donors *×*3 activation states: resting, 8hrs post-stimulation, and 48hrs post-stimulation). Unlike the immortalized lines in Replogle22/Nadig25, this dataset captures biologically relevant heterogeneity from both activation state and inter-individual variation [Brodin et al., 2015, Chapfuwa et al., 2025].

#### Context-aware splitting strategy

To fairly compare *within-context* and *cross-context* generalization, we design a splitting strategy that aligns the two evaluation regimes on three axes: held-out perturbations, test-set size, and training-set size (Figure 7, Appendix). Both splits are derived from a shared random seed and planning step to ensure identical randomization decisions, see Appendix C.2 for details.

**Figure 7:**
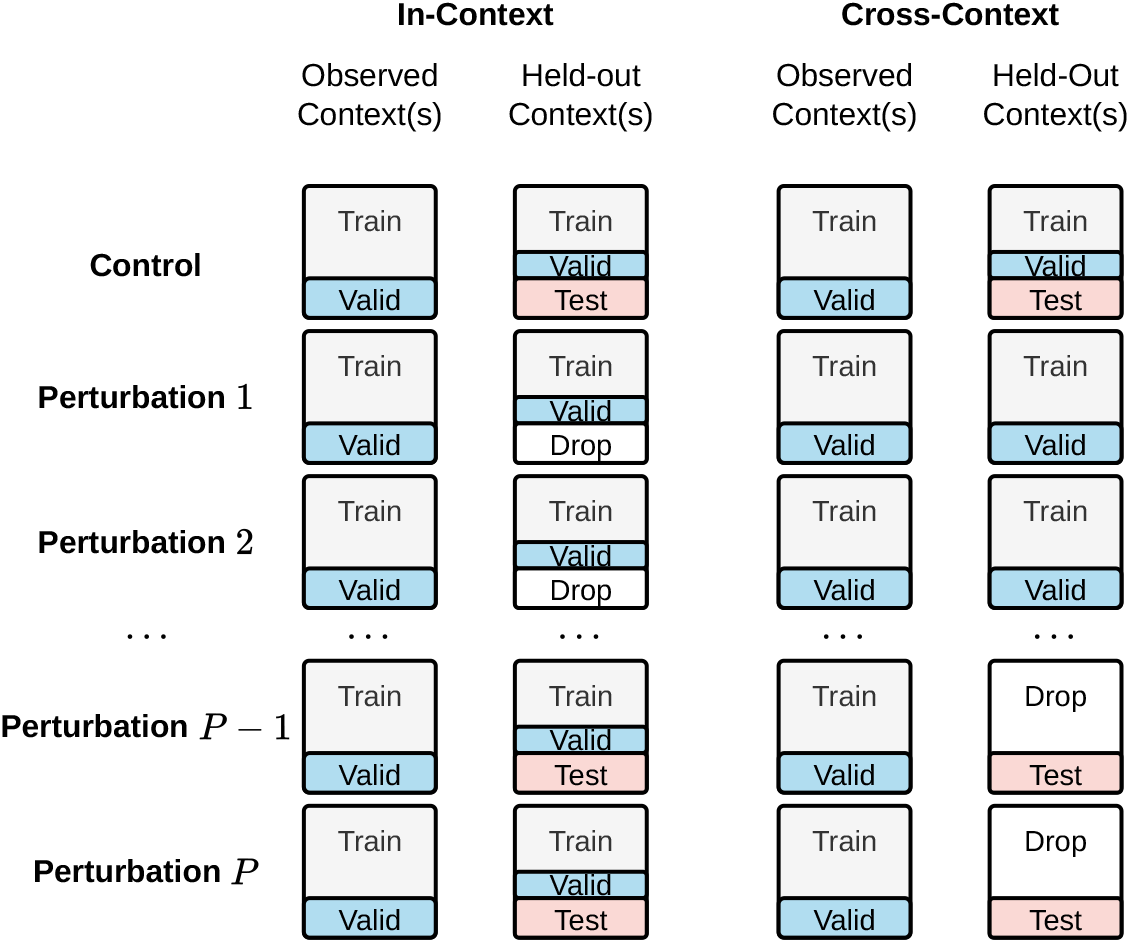
Within-context vs. cross-context splitting strategy. Cells marked “Drop” are removed during fairness alignment.

### 4.3 Results

#### 4.3.1 Validation and robustness of the Vendi score

We compare Vendi score with covariance effective rank [Roy and Vetterli, 2007], the exponential entropy of the normalized covariance eigenvalues, to assess the value of Vendi’s null-calibrated distance kernel. Across 100 synthetic DIRECTDGP datasets, both measures increase with the number of perturbations *P* (Figure 8, Appendix E.2). However, Vendi tracks increasing effect strength from the population null, whereas effective rank remains high under weak effects (Figure 3a). Moreover, sample-size and deterministic-prediction biases are largest at low effect factors and diminish with increasing effect strength and larger *N*_*q*_ (Figure 10, Appendix). In Gaussian-noise experiments on NORMAN19 (Figure 9, Appendix E.2), Vendi and PDS (*l*_1_) remain stable under low-to-moderate noise and decline under severe noise, whereas effective rank increases with noise.

**Figure 8:**
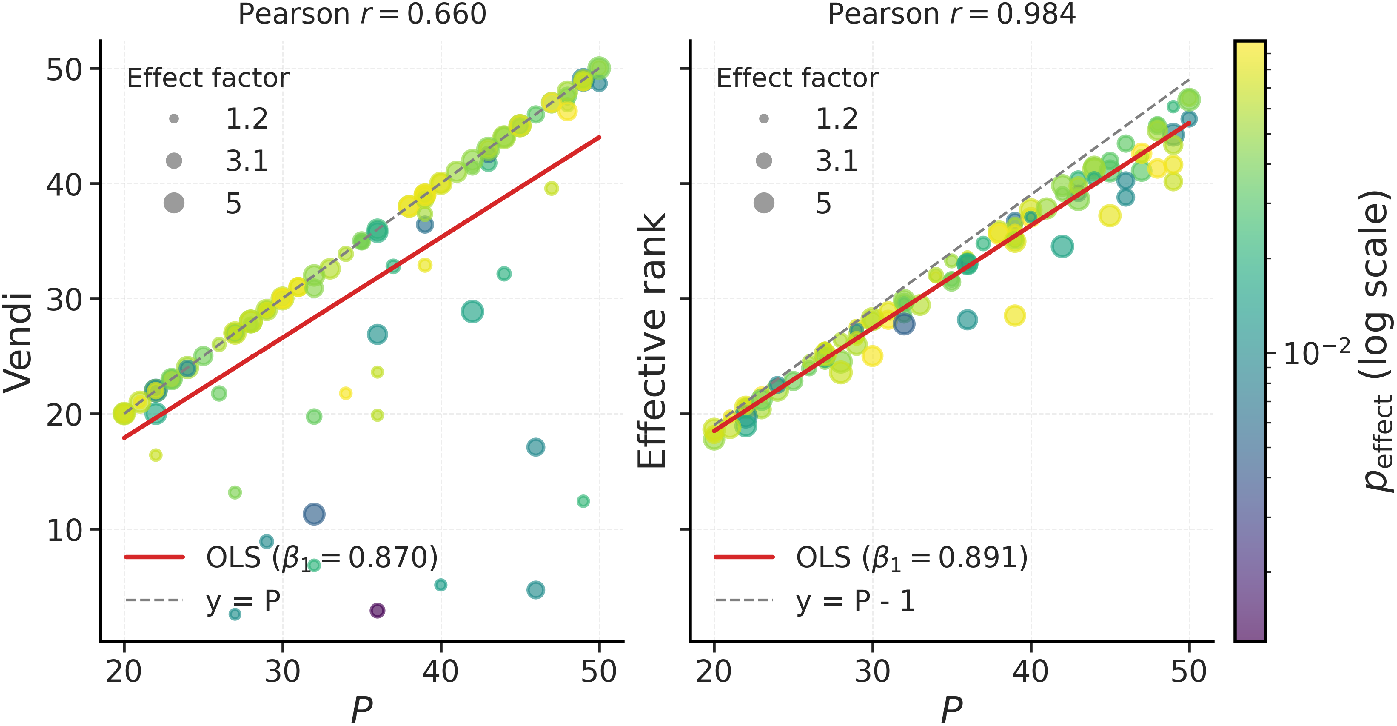
Raw empirical Vendi score (left) and covariance effective rank (right) versus the number of perturbations *P* across 100 randomly sampled synthetic DirectDGP datasets. Each circle is one dataset. Color encodes the perturbation-effect probability *p*_effect_ on a logarithmic scale, and marker size encodes the effect factor. Red lines show ordinary least-squares fits, with the estimated slopes reported in the legends; Pearson correlations are reported above each panel. Gray dashed lines denote the reference relationships VS(*Y* ) = *P* and erank(*Y* ) = *P* − 1, respectively.

**Figure 9:**
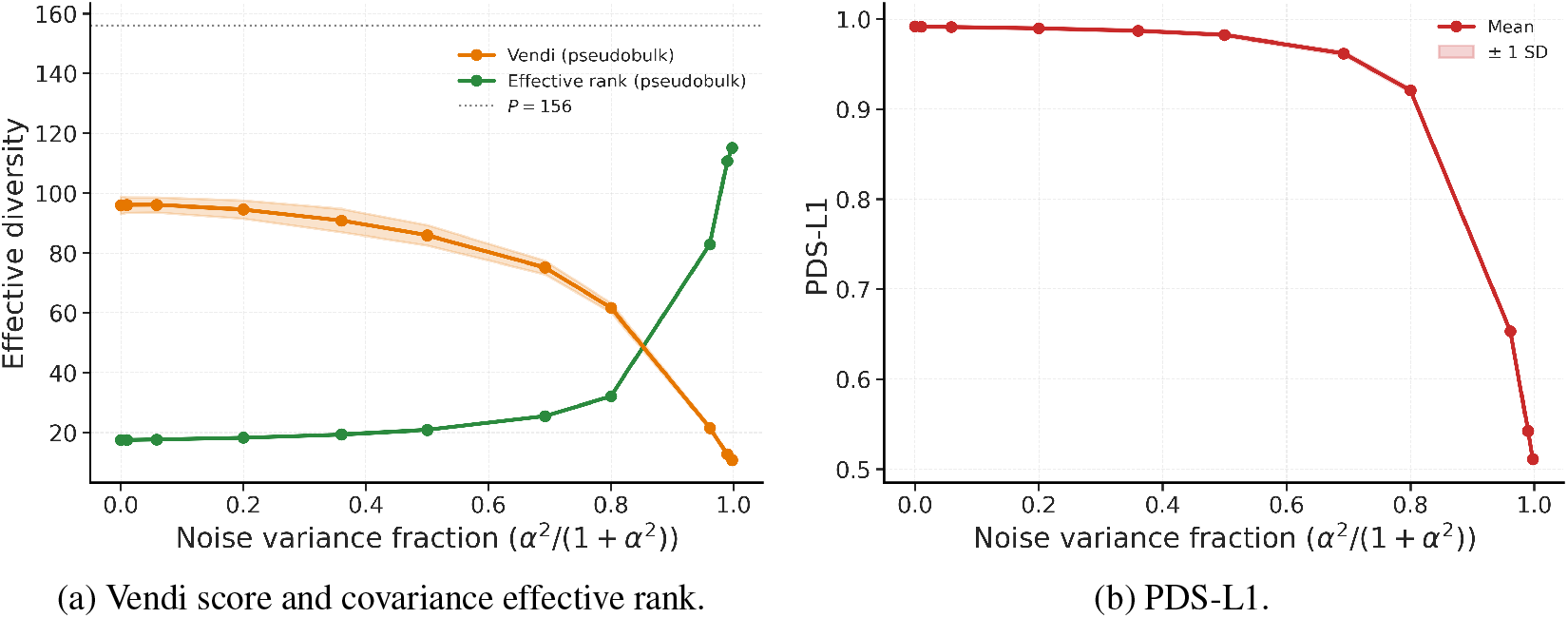
Robustness to Gaussian noise on the Norman19 dataset. We inject Gaussian noise scaled to each gene’s own variability, *σ*_*g*_ = *α* std_*g*_, so *α* is a dimensionless noise-to-signal ratio and the noise variance fraction is *α*^2^*/*(1 + *α*^2^). (a) Vendi score and covariance effective rank as functions of the noise variance fraction. (b) PDS (*l*_1_) over the same noise sweep. Lines and markers show the mean across 10 noise realizations, and shaded bands indicate *±*1 standard deviation.

**Figure 10:**
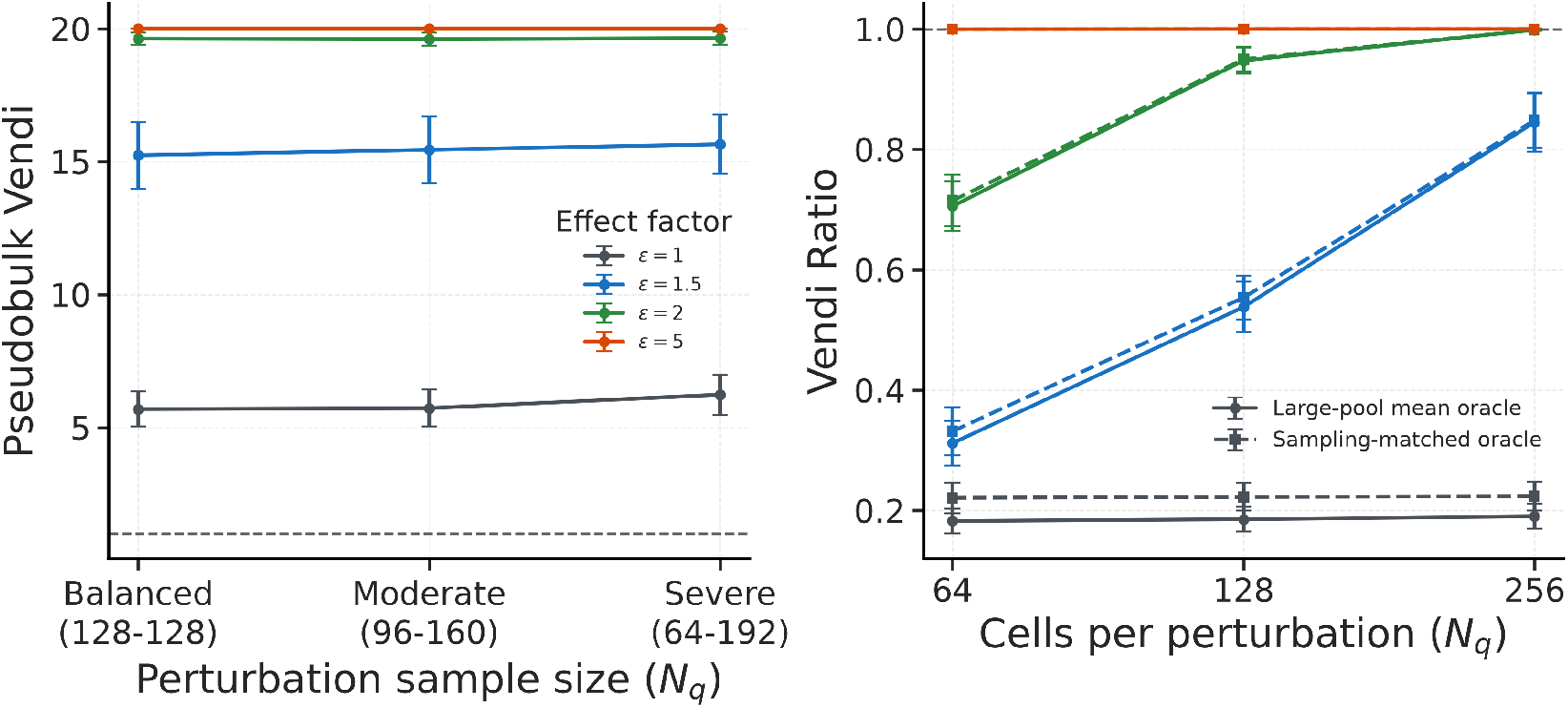
Sensitivity of Vendi score to perturbation sample size and finite-sample noise. (left) Mean Vendi score across balanced (*N*_*q*_ = 128), moderately heterogeneous (*N*_*q*_ = 96–160), and severely heterogeneous (*N*_*q*_ = 64–192) sampling approach. All approaches have the same median *N*_*q*_ = 128 and total number of perturbed cells. Colors denote DirectDGP effect factors *ϵ* ∈ {1, 1.5, 2, 5} . (right) Vendi Ratios for a large-pool mean oracle, computed from 2,048 cells per perturbation, and an independent sampling-matched oracle containing the same *N*_*q*_ as the empirical target. Ratios are shown for *N*_*q*_ ∈ {64, 128, 256} ; colors denote effect factor, while solid circles and dashed squares denote the large-pool and sampling-matched oracles, respectively. Points and error bars show mean *±* one standard deviation across 10 paired simulation seeds. Simulations used *P* = 20, *G* = 1024, *N*_0_ = 512, *p*_effect_ = 0.1, and *β* = 0. Horizontal dotted lines indicate a Vendi score or Vendi Ratio of one.

#### 4.3.2 Perturbation response geometry: diversity and discrimination

We characterize perturbation-response geometry through relative diversity (VR; (8)) and discrimination (PDS; (13); Figure 3b). On NORMAN19, most models achieve high PDS but low VR, suggesting partial diversity collapse; scVI is well calibrated, whereas scLDM overestimates diversity. Across datasets, scLDM performs among the strongest models, leading on Replogle22/Nadig25, while STATE-Geneformer attains the highest PDS on Norman19. All models underestimate diversity on Zhu25, where GEARS also exhibits near-random PDS. On Replogle22/Nadig25 (RPE1–K562), the largest perturbation set, PDS is lower than on Norman19, although scLDM retains strong discrimination and near-unity VR. Notably, under cross-context evaluation, PDS broadly declines, approaching random performance on ZHU25 (Figure 11; Appendix). Thus, unseen perturbation–context combinations consistently impair discrimination but affect diversity in a model- and dataset-dependent manner.

**Figure 11:**
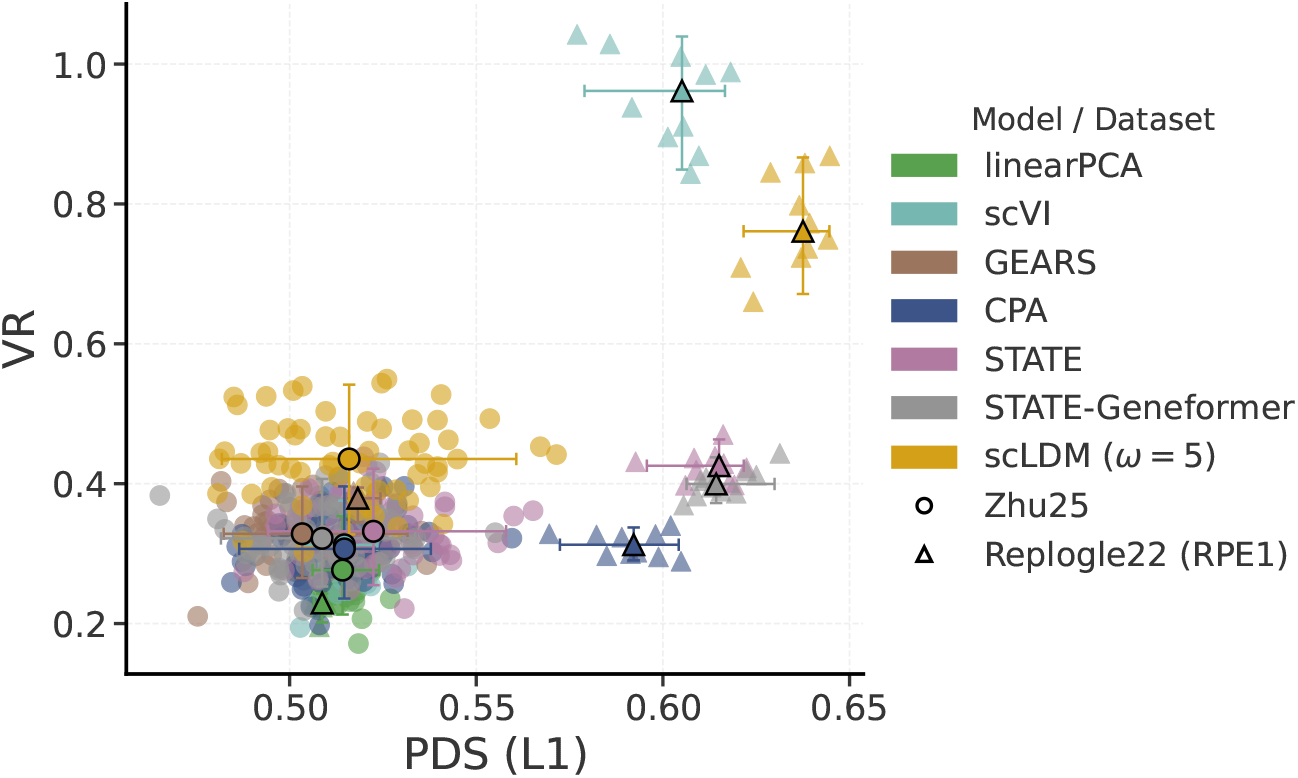
Vendi ratio (VR) vs. PDS (*l*_1_) across models and datasets under cross-context evaluation. Color encodes the model and marker shape encodes dataset. Bold markers and error bars show the median and 95% percentile intervals per model–dataset pair. Higher PDS indicates better discrimination; VR = 1 indicates agreement with empirical effective diversity.

#### 4.3.3 Cross-context generalization gap

Figure 4 summarizes the cross-context generalization gap on CausalDGP and Zhu25 (full results in Figures 12–15, Appendix). On CausalDGP, scLDM and CPA outperform the baselines within context. Cross-context performance remains high when only the baseline drift *B*_*C*_ changes, consistent with Proposition 1, under which latent-effect transportability is independent of *B*_*C*_. Performance falls to baseline levels when perturbation-relevant mechanisms in *A*_*C*_ change, alone or together with *B*_*C*_, consistent with a loss of direct latent transportability.

**Figure 12:**
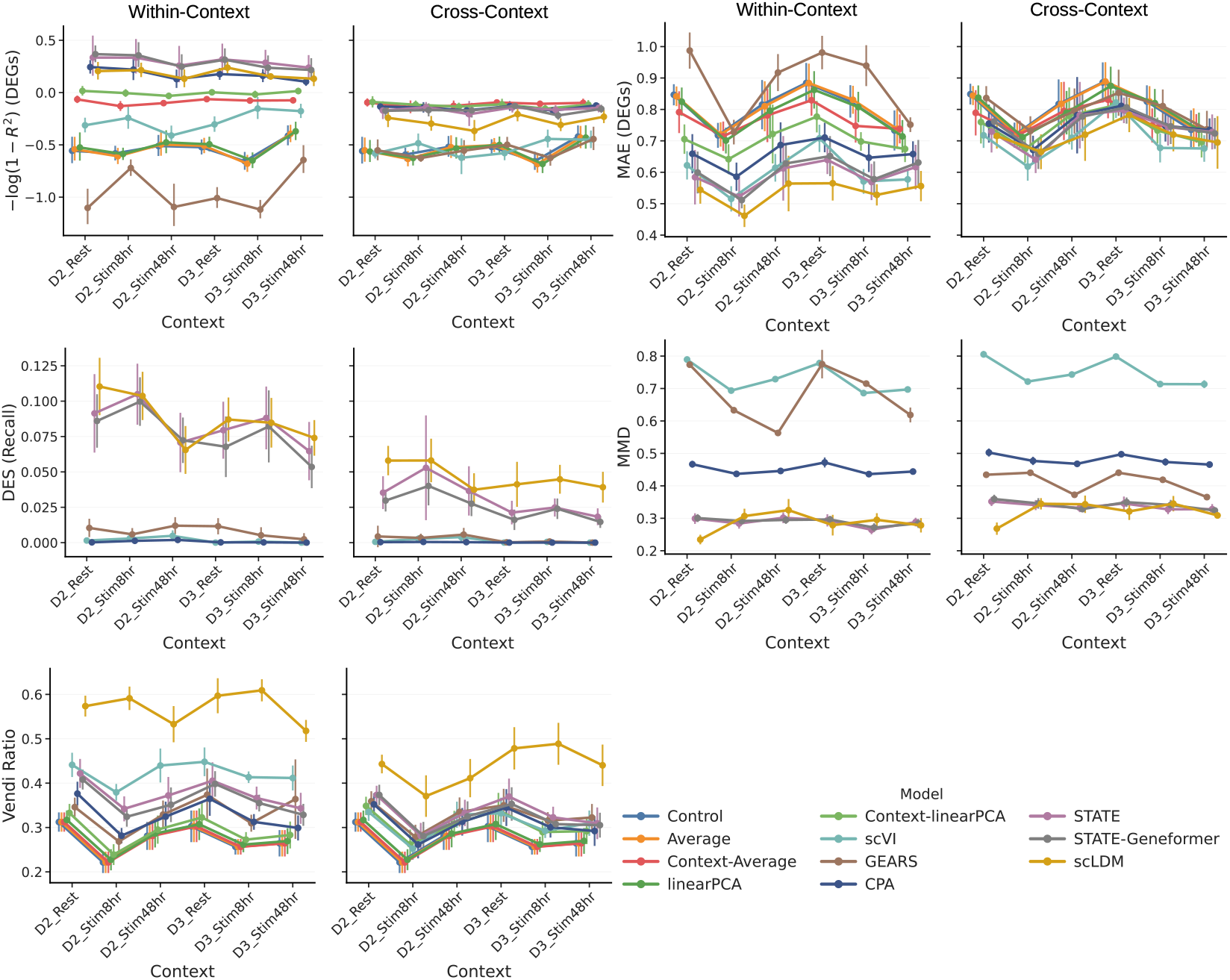
Additional results on the cross-context generalization gap in Zhu25 under within-context and cross-context evaluation. The *x*-axis denotes the six test contexts (2 held-out donors *×* 3 activation states). We plot medians and interquartile ranges. scLDM is evaluated with *ω* = 5.

On Zhu25, performance similarly declines across the held-out donors and activation states. While CPA, STATE, scLDM, and STATE-Geneformer outperform simple baselines within-context evaluation, their advantage largely vanishes in cross-context splits. This suggests donor-specific variation in the causal mechanisms governing CD4+ T-cell perturbation responses, potentially driven by environmental factors, pathogen exposure history, or inter-individual genetic variation [Brodin et al., 2015, Chapfuwa et al., 2025].

On Replogle22/Nadig25 (Figure 5; full results in Figure 13–14, Appendix E.4), where models are trained on RPE1 and evaluated on held-out K562 perturbations, the cross-context gap is notably smaller: CPA, STATE, and STATE-Geneformer remain competitive with simple baselines even under cross-context evaluation. Despite weak overall correlation between these cell lines [Nadig et al., 2025], conserved effects for some perturbations may support transport of the transferable signal.

**Figure 13:**
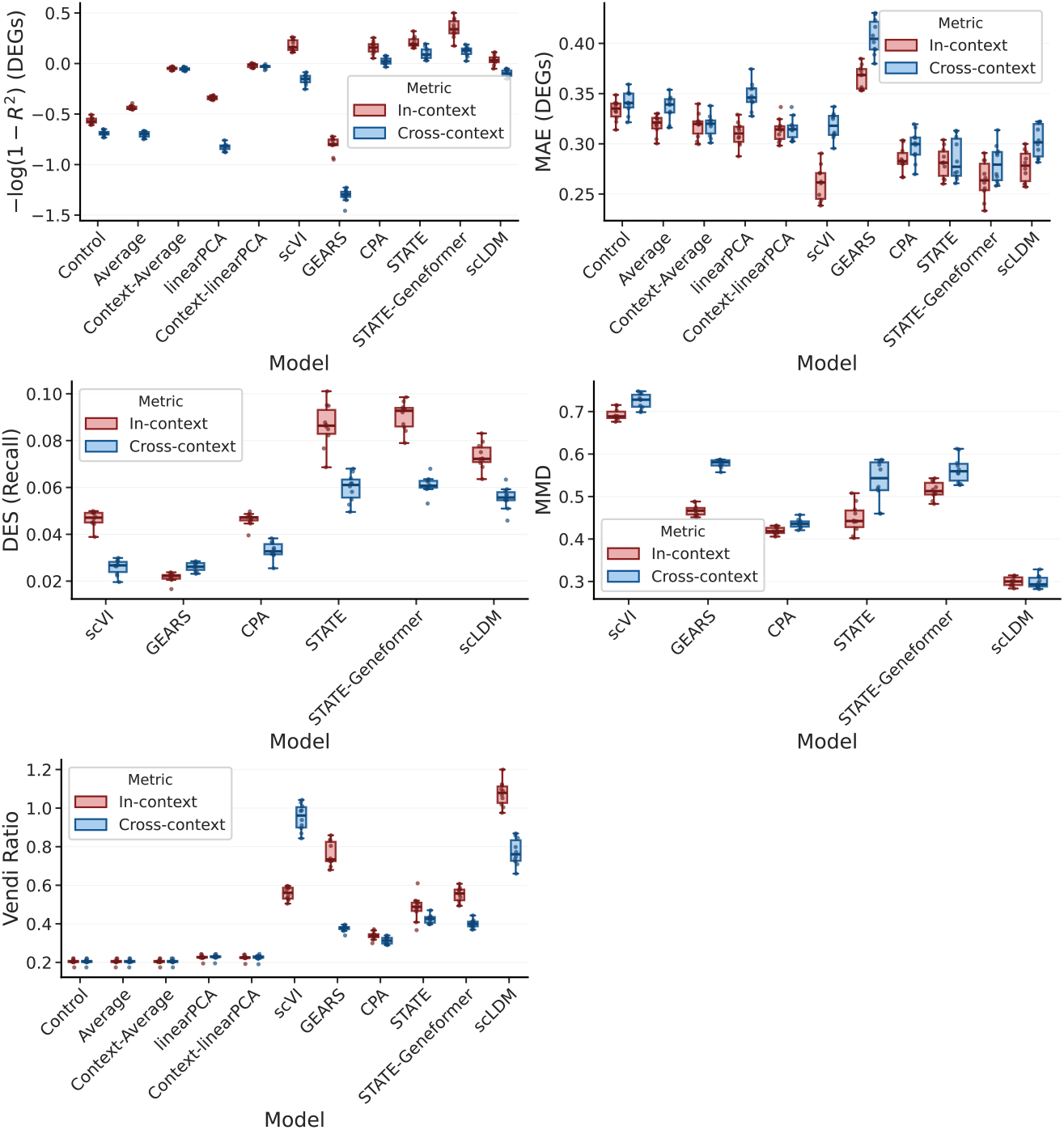
Additional results for the cross-context generalization gap on Replogle22/Nadig25 (RPE1–K562) under within-context vs. cross-context evaluation. Models are trained on RPE1 and evaluated on held-out K562 perturbations. scLDM is evaluated with *ω* = 5.

##### Alignment between theory and experiments

We test the prediction in (6) that direct-transport performance declines as *perturbation-relevant causal mechanisms* diverge across contexts, using both real Perturb-seq datasets and CausalDGP. As a proxy for mechanistic similarity, we use the Pearson correlation between source- and target-context noise-adjusted TI values across matched perturbations; TI is defined in (19) [Nadig et al., 2025]. Across three Replogle22/Nadig25 source-to-K562 tasks, we evaluate the four best-performing models: CPA, STATE, STATE-Geneformer, and scLDM, each trained separately on RPE1, Jurkat, and HepG2. Jurkat and HepG2 exhibit similarly high TI concordance with K562, whereas RPE1 exhibits lower concordance, which is biologically plausible given RPE1’s non-cancer origin (Table 2). Consistent with this difference, RPE1 → K562 yields the largest transfer gaps across all four models for both DEG-restricted Pearson ***δ*** and PDS (*l*_1_) (Figure 14, Appendix). Moreover, varying *A* in CausalDGP reproduces the observed TI–transfer-gap relationship across the Replogle22/Nadig25 TI range. Cross-donor TI correlations in Zhu25 are substantially lower and coincide with broad generalization failures (Figure 4). Although TI correlation is only a proxy and does not identify the underlying causal mechanisms, consistency across models and across real and synthetic Perturb-seq datasets supports the predicted association between greater mechanistic divergence and reduced direct-transport performance.

**Table 2:**
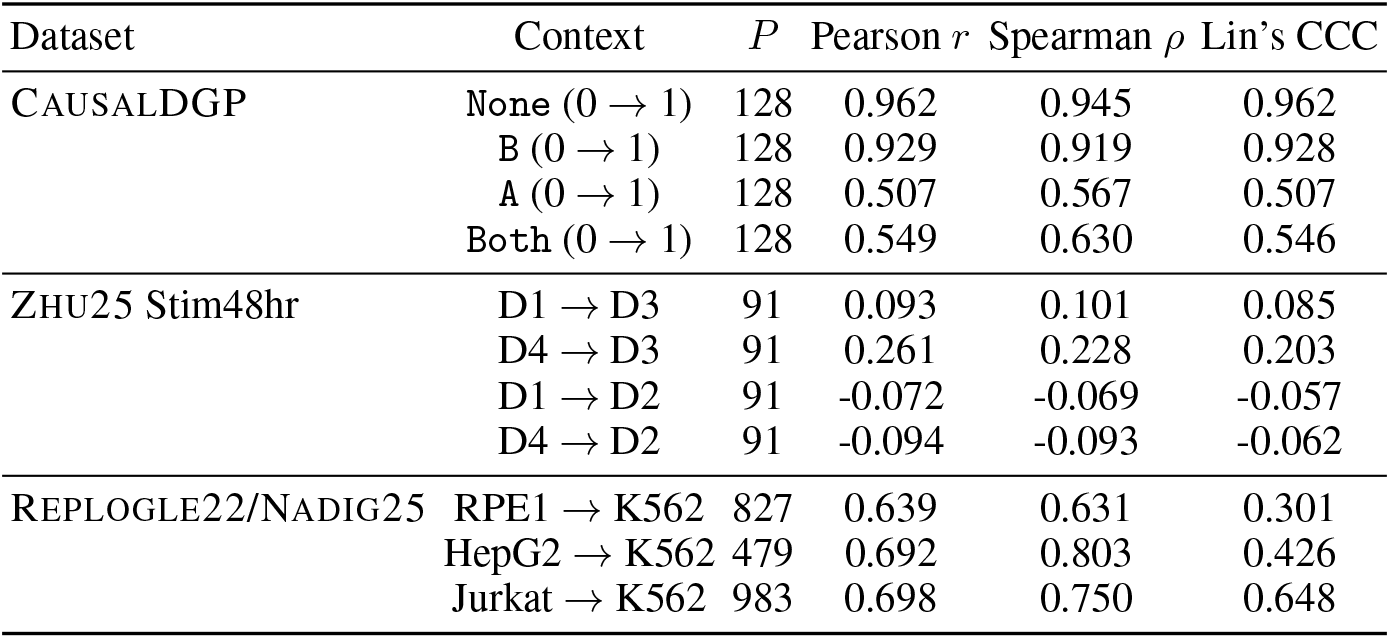
Cross-context concordance of TI. For each dataset and context pair, we compare per-perturbation TI estimates between the shared *P* perturbations in the source and target contexts using Pearson *r*, Spearman *ρ*, and Lin’s CCC [Lin, 1989], which penalizes deviation from *y* = *x*.

**Figure 14:**
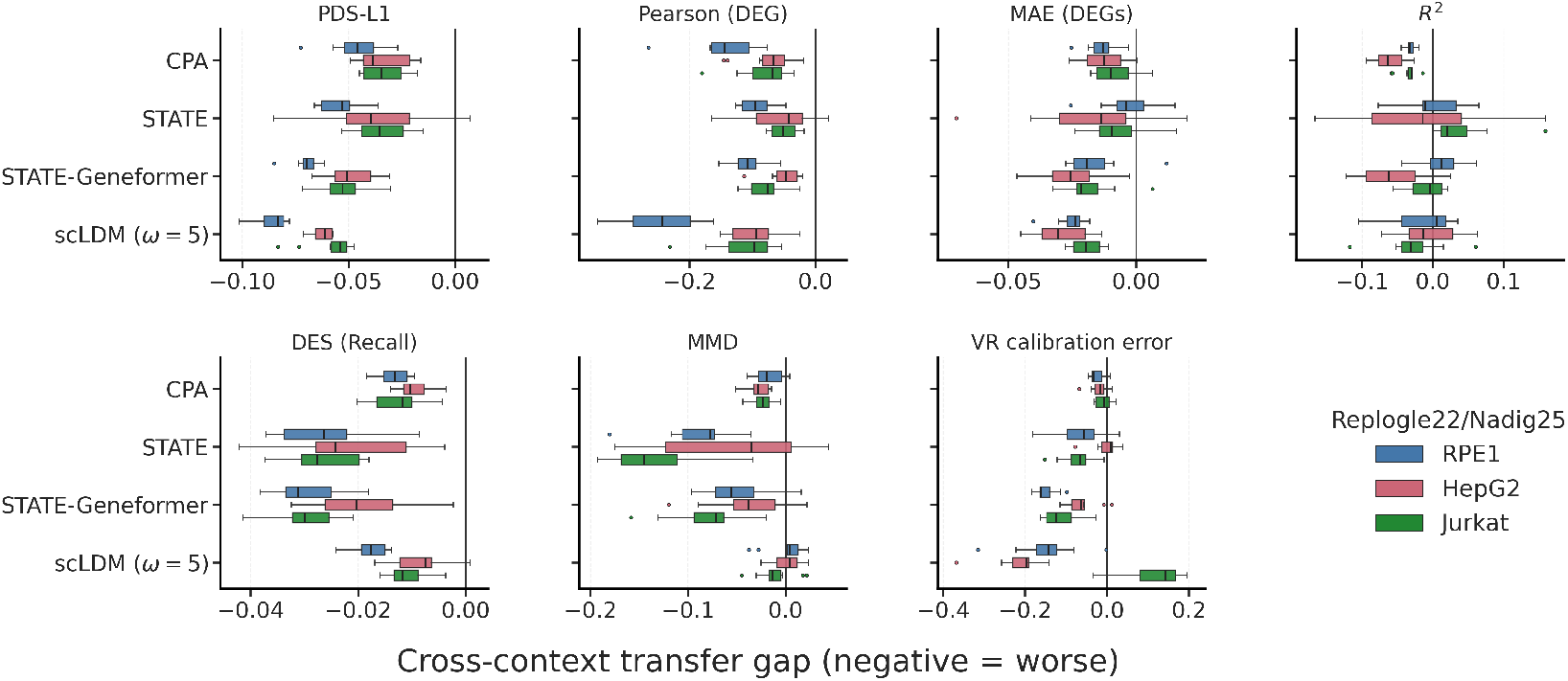
Cross-context transfer gaps on Replogle22/Nadig25 for RPE1–K562, HepG2–K562, and Jurkat–K562. Models are trained in the indicated source context and evaluated on held-out K562 perturbations. Box plots summarize 10 paired trial-level gaps. Gaps are oriented so that negative values consistently indicate worse cross-context performance. For VR, the gap is the cross-context decrease in absolute deviation from perfect calibration (VR = 1), so negative values indicate worse calibration.

## 5 Conclusion

Cross-context generalization in perturbation prediction depends on shared causal mechanisms, not merely distributional similarity. Across six deep learning models and five simple baselines, performance drops substantially under cross-context evaluation, often matching simple baselines. A joint Vendi–PDS analysis characterizes perturbation-response geometry along effective-diversity and discrimination axes, revealing complementary failure modes: simple baselines produce low-diversity, poorly differentiated responses, whereas deep learning models generally achieve stronger in-context discrimination but often underestimate diversity and consistently lose discrimination under cross-context transfer. Even on synthetic data with known causal structure, no model generalizes across contexts with divergent mechanisms, exposing limitations of current architectures. Consistent with theory, transfer gaps widen as cross-context transcriptome-wide impact (TI) concordance declines, supporting low TI concordance as an empirical proxy for mechanistic divergence.

## Acknowledgments and Disclosure of Funding

We thank Julia Greissl, Jabran Zahid, Christopher Gooley, Lorin Crawford, and Miah Wander for fruitful discussions. Shi-ang Qi conducted this work during an internship at Microsoft Research.

## A Baselines

Table 3 summarizes the considered baselines. Control predicts the mean expression of all training control cells, whereas Average predicts the cell-weighted mean across all non-control training cells; both return the same vector for every perturbation and context. Table 4 presents detailed transport formulae for the remaining models. The CausalDGP simulator shares inductive biases with these methods but realizes them differently; we therefore group methods by their primary assumptions rather than their architectures. *Transport-based* methods learn a map from control to perturbed distributions: CausalDGP uses a linear SDE, scLDM learns a neural velocity field through conditional flow matching, and STATE uses optimal-transport-based matching between control and perturbed populations. STATE-Geneformer retains the STATE transport mechanism but represents basal cell states using pretrained Geneformer embeddings. *Disentanglement-based* methods (CPA/scVI) decompose latent variation into perturbation, context, and cell-state components. Finally, GEARS encodes explicit gene–gene structure using Gene Ontology and basal co-expression graphs.

**Table 3:** Perturbation prediction models and their transport assumptions for estimating *P* (*Y* | do(*Q*=*q*), *C*=*c*_test_). Full transport formulae are given in Table 4 (Appendix A).

| Model | Transport mechanism | Context-aware | Biological prior | Granularity |
| --- | --- | --- | --- | --- |
| Control | Control mean, identity (no perturbation signal) | ✗ | ✗ | Pseudobulk |
| Average | Perturbation average | ✗ | ✗ | Pseudobulk |
| Context-Average | Context-specific perturbation average | ✓ | ✗ | Pseudobulk |
| linearPCA | Additive in gene space | ✗ | ✗ | Pseudobulk |
| Context-linearPCA | Context-specific, additive in gene space | ✓ | ✗ | Pseudobulk |
| scVI | Additive in latent space | ✓ | ✗ | Single-cell |
| CPA | Additive in latent space | ✓ | ✗ | Single-cell |
| GEARS | Additive residual in gene space | ✗ | Gene Ontology and co-expression graphs | Single-cell |
| STATE | Nonlinear (attention) | ✓ | ✗ | Single-cell |
| STATE-Geneformer | Nonlinear (attention) | ✓ | Pretrained Geneformer embedding | Single-cell |
| scLDM | Conditional flow matching in VAE latent space | ✓ | ✗ | Single-cell |

**Table 4:**
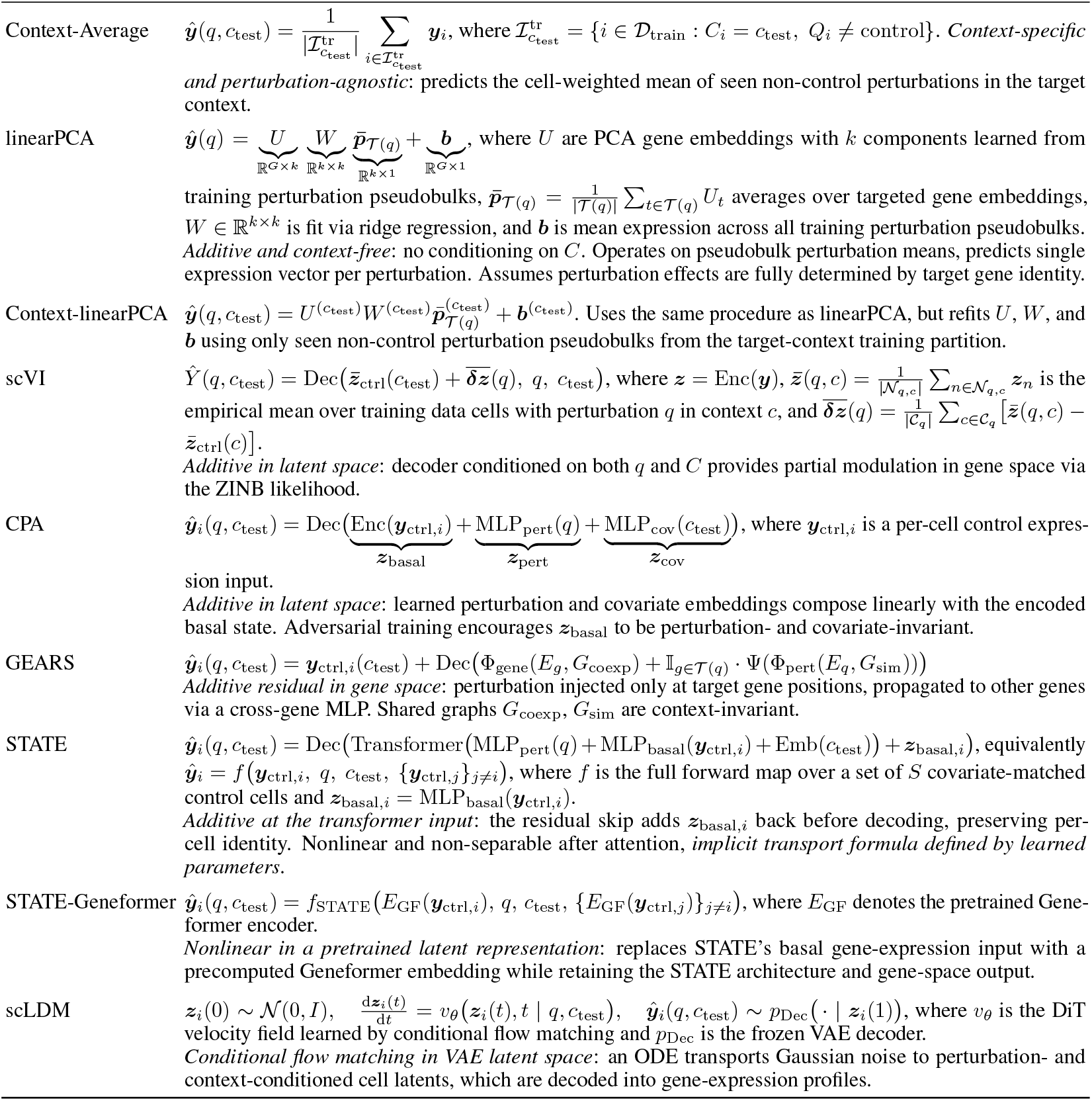
Perturbation prediction models and their assumed transport formulae for estimating *P* (*Y* | do(*Q* = *q*), *C* = *c*_test_) from training data {(*q, c*_train_), (*q*^*′*^, *c*_test_) : *q*^*′*^*≠ q*}.

## B Data generation process

Here we provide additional implementation details for the two data-generating processes used in this paper. See Table 5 for a summary of the hyperparameters.

**Table 5:** Simulation parameters for the two semi-synthetic data-generating processes used in this work. DirectDGP, adapted from Mejia et al. [2025], is primarily used for within-context metric stress tests, whereas CausalDGP is the proposed two-context mechanistic simulator for studying transportability under controlled mechanism shift.

| Parameter | Meaning | DIRECTDGP | CAUSALDGP |
| --- | --- | --- | --- |
| *Shared parameters* |  |  |  |
| $\vert C\vert $ | Number of biological contexts | 1 | 2 |
| $G$ | Number of genes | 1024–8192 | 128 |
| $N_0$ | Number of control cells | 128–1024 | 1024 |
| $N_q$ | Number of cells under perturbation $q$ | 128–512 | 1024 |
| $P$ | Number of perturbations | 20–50 | 128 |
| $\theta_g$ | Gene-wise inverse-dispersion | Estimated from NORMAN19 | Estimated from NORMAN19 |
| *DirectDGP-specific parameters* |  |  |  |
| $p_{\text{effect}}$ | Probability that a gene is perturbed | 0.001–0.1 | - |
| $\epsilon$ | Multiplicative perturbation factor | 1.2–5.0 | - |
| $\beta$ | Control-bias scale | 0.0–2.0 | - |
| $\mu_\ell$ | Mean log-library size | 0.2–5.0 | - |
| $\sigma_\ell$ | standard deviation of log-library size | 0.1714 | - |
| $\mu_g^c$ | Baseline control mean | Estimated from NORMAN19 | - |
| $\lambda_g$ | Gene-specific control-bias term | Estimated from NORMAN19 | - |
| *CausalDGP-specific parameters* |  |  |  |
| $A_c$ | Context-specific interaction matrix | - | Sparse, randomized, spectrally shifted to be Hurwitz |
| $B_c$ | Context-specific baseline bias | - | Entries sampled from $\text{Unif}([-3, 3])$ |
| $\Gamma_q$ | Perturbation shift vector | - | One-sparse intervention |
| $s_q$ | Perturbation magnitude | - | $\text{Unif}([-6 \log G, -2 \log G] \cup [2 \log G, 6 \log G])$ |
| $\rho_{\text{swap}}$ | Row-wise rewiring fraction | - | 0.5 |
| $\sigma$ | Diffusion coefficient in latent OU process | - | $\sqrt{2}$ |
| $\beta_{\text{sp}}$ | Softplus parameter | - | 3 |

### B.1 Additional details for DirectDGP

DirectDGP is adapted from the simulation framework of Mejia et al. [2025]. We refer the reader to that work for the full derivation and motivation, and depict the process in Algorithm 2 for completeness. Briefly, DirectDGP generates single-cell perturbation data in a *single biological context*, with gene-wise negative-binomial counts calibrated to mimic key statistical properties of Norman19, including sparse counts, heterogeneous library size, and control-referenced bias. In this paper, we use DirectDGP primarily as a controlled within-context benchmark for studying metric behavior, including metric distortion and Vendi score validation.

### B.2 Additional details of CausalDGP

This section provides additional implementation details for CausalDGP. Recall from Section 2.2 that, conditional on biological context *C* ∈ {0, 1} and perturbation *Q* ∈ {0, 1, …, *P*}, the latent state *X*(*t*) ∈ ℝ^*G*^ evolves according to an affine OU process, and the observed counts are generated through a gene-wise negative binomial emission process. Here we specify how the simulator instantiates the causal interaction matrix *A*_*C*_, the context-specific baseline drift *B*_*C*_, the shift vector Γ_*Q*_, and the sampling procedure.

#### Gene sub-sampling and empirical anchoring

To anchor the observation model to realistic single-cell count statistics, we subsample *G* genes without replacement from a larger empirical pool of *M* genes whose dispersion parameters have been estimated from the Norman19 dataset:

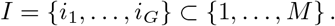

We retain the corresponding gene names and inverse-dispersion parameters,

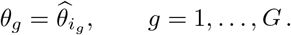

#### Construction of the sparse causal interaction matrix

As we described in Section 2.2, the entry convention for the causal interaction matrix *A*_*C*_ is

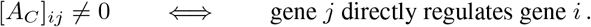

Thus, the deterministic drift of gene *i* depends on its upstream regulators:

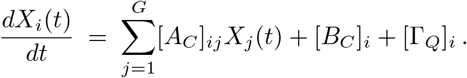

For each target gene *i*, we sample exactly 10 distinct regulators from {1, …, *G*}*\*{*i*} and assign each edge an independent signed weight from Unif [−3, −1] ∪ [1, 3] . We use this fixed-in-degree construction to control network density and local regulatory complexity, thereby isolating context-specific mechanism shifts from variation in in-degree. This choice may underrepresent heterogeneity in perturbation propagation arising from highly connected regulators in biological networks. Moreover, a randomly generated sparse matrix is not necessarily stable. To guarantee that (2) admits a unique stationary Gaussian distribution, we stabilize each raw matrix by shifting it by a multiple of the identity:

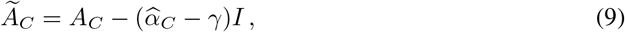

where 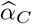 estimates the spectra abscissa (largest real part among eigenvalues) of the raw *A*_*C*_ and *γ <* 0 is a user-chosen stability margin (set to *γ* = −1 throughout). If *λ* is an eigenvalue of *A*_*C*_, the corresponding eigenvalue of *Ã*_*C*_ is

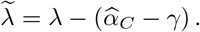

When the spectral–abscissa estimate is accurate,

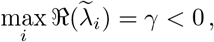

so *Ã*_*C*_ is Hurwitz. Throughout the paper we overload notation and write *A*_*C*_ for the stabilized matrix.

#### Mechanism-level context diversity

The two contexts *C* ∈ {0, 1} differ through their latent mechanisms (*A*_*C*_, *B*_*C*_), not through post-hoc output perturbations. We define four diversity modes:

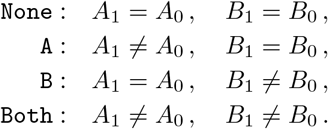

When matrix diversity is enabled, *A*_1_ is obtained by row-wise rewiring of *A*_0_: in each row a fraction of existing nonzero entries is moved to previously zero positions, preserving both the row sparsity pattern and the multiset of edge weights while changing which regulators act on each target. Each context matrix is then stabilized independently via the identity shift in (9), since rewiring can alter the spectral abscissa. When bias diversity is enabled, *B*_1_ is resampled coordinate-wise from [*B*_1_]_*g*_ ∼ Unif([*−*3, 3]). By construction, cross-context non-transportability arises directly from mechanism shift.

##### Algorithm 2

DirectDGP

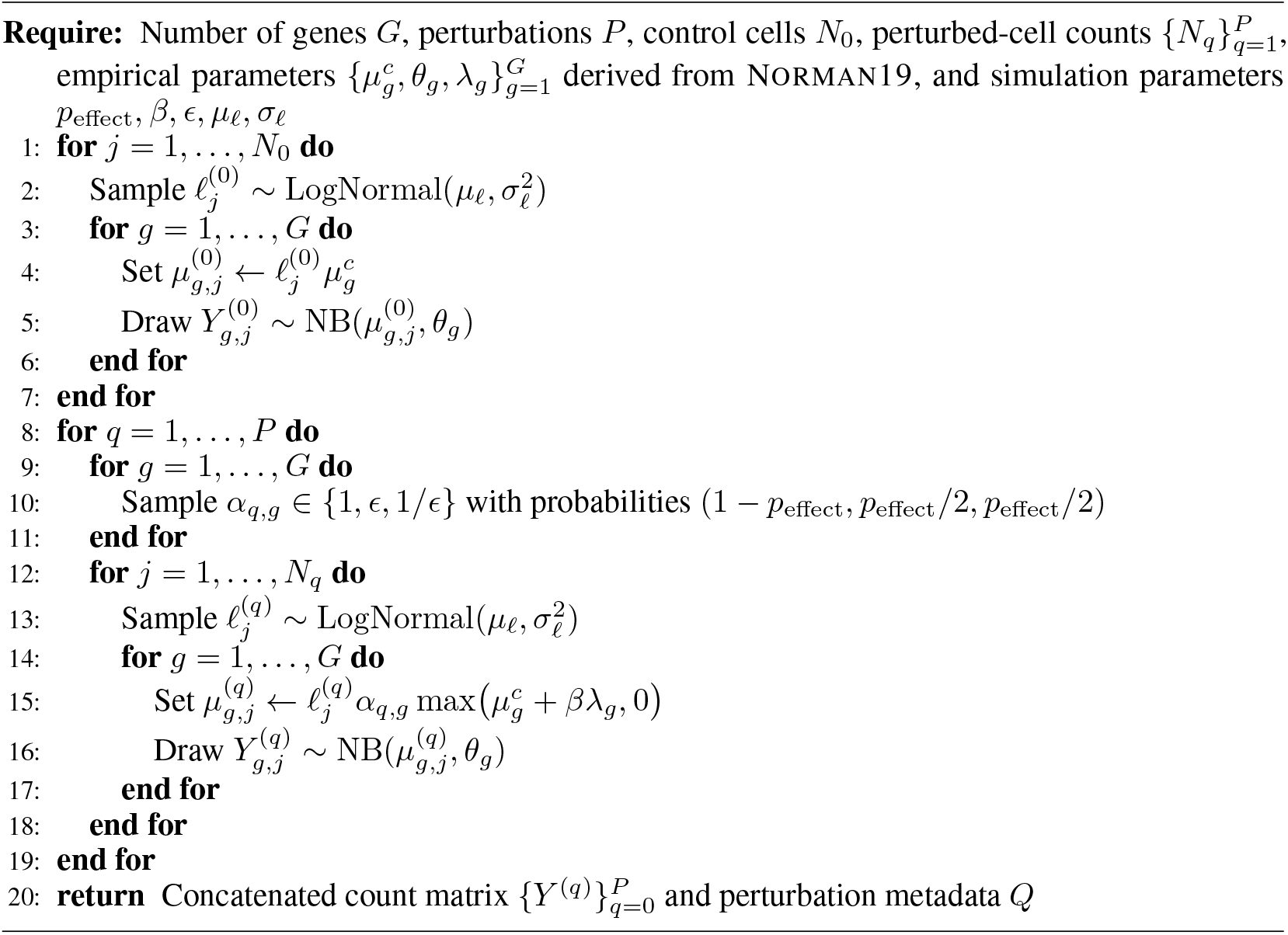

#### Perturbation shift vector

Each non-control perturbation targets exactly one gene. We sample *P* distinct targets

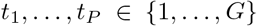

without replacement and define

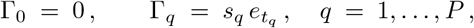

where 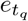 is the *t*_*q*_-th standard basis vector and the perturbation magnitudes *s*_*q*_ are drawn from

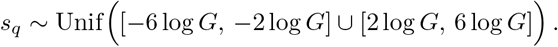

Each perturbation therefore acts directly on a single coordinate but propagates globally through the network dynamics.

#### Euler–Maruyama sampling

Although the stationary distribution of (2) is Gaussian in latent space, we sample from it via Euler–Maruyama (EM) discretization with step size Δ*t*:

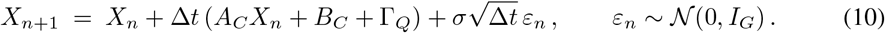

For each context–perturbation pair, we simulate up to 64 parallel trajectories, independently initialized from *N*(0, *I*_*G*_) and driven by independent Gaussian increments. We use Δ*t* = 10^−3^, burn in each trajectory for 12,000 steps, and collect one state per trajectory per sampling round, advancing each trajectory by 10 steps between rounds. Conditional on the simulator parameters, different trajectories are independent. Let

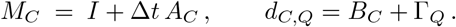

Then (10) becomes

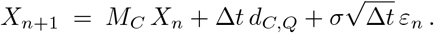

#### Emission process and count parameterization

Given a latent state ***x* ∈** ℝ^*G*^, we map it to nonnegative gene-wise mean expression via the softplus link:

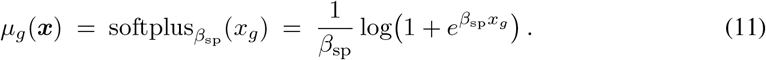

The observed count for gene *g* is then drawn independently as

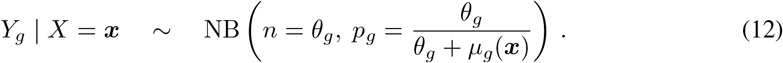

Under this parameterization,

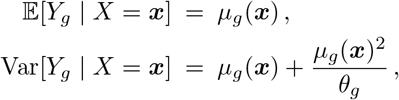

recovering the standard negative-binomial mean–dispersion model used in scRNA-seq analysis.

#### Condition-wise sampling scheme

Within each condition, context labels are assigned independently as

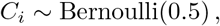

so the realized number of cells per context is random rather than exactly balanced. Conditional on (*C*_*i*_, *Q*_*i*_), we instantiate the corresponding EM sampler, draw a latent state, transform it via the softplus link, and sample observed counts from the negative-binomial model. Algorithm 3 summarizes the full procedure for generating the CausalDGP dataset.

##### Algorithm 3

CausalDGP

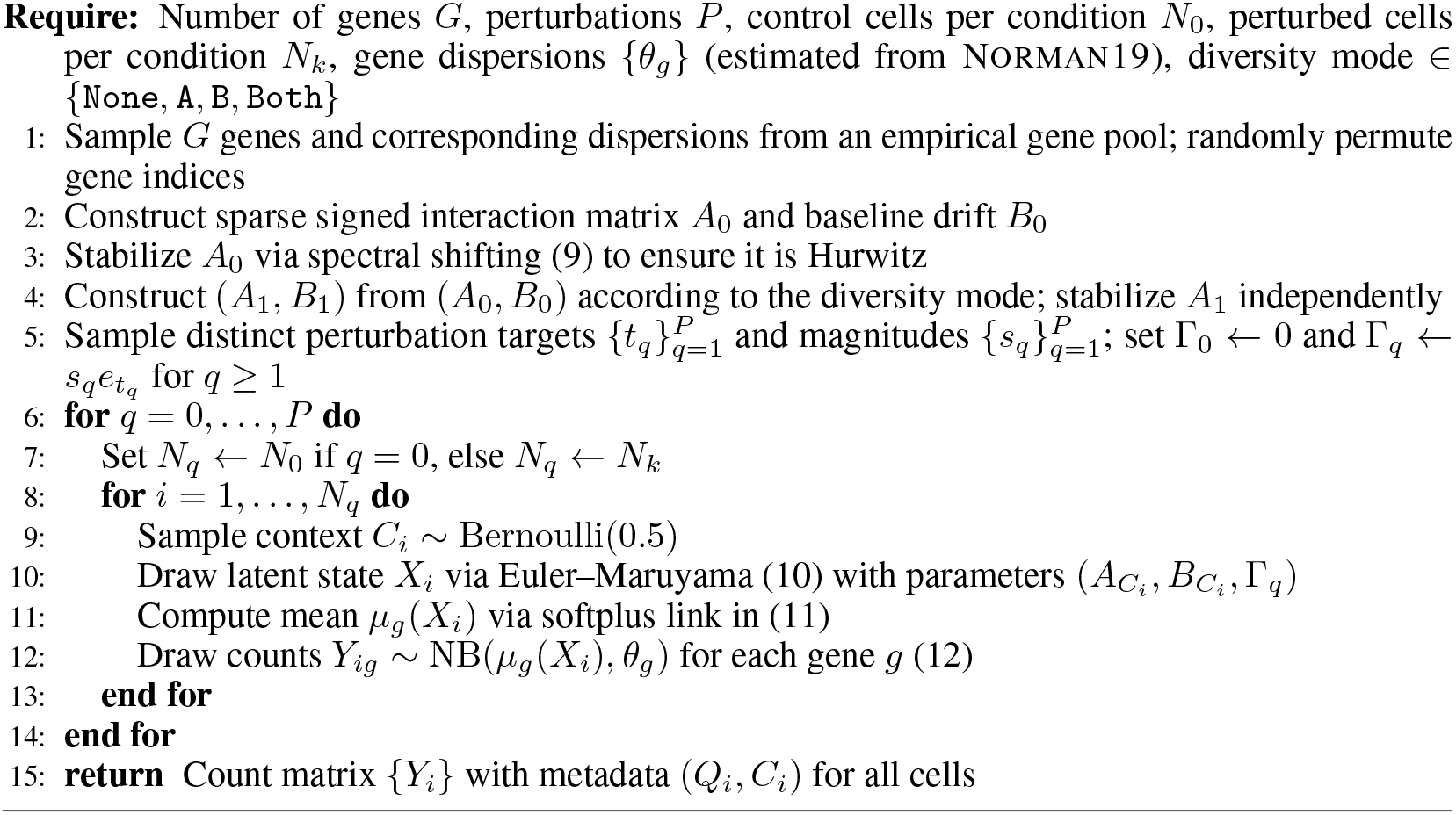

#### Remarks

Two implementation details are worth noting. First, because the emission process is shared across contexts, all non-transportability in CausalDGP arises from latent mechanism differences in (*A*_*C*_, *B*_*C*_), not from measurement-level shift. Second, because the affected-gene masks are defined from equilibrium mean shifts, they provide exact ground truth for evaluating whether a model recovers the downstream support of each perturbation effect.

#### B.2.1 Validating CausalDGP

##### Qualitative cell-level heterogeneity

Figure 6 illustrates how CausalDGP generates context variation at the mechanism level and how this variation manifests in the simulated single-cell expression manifold. The heatmaps in the top row display the latent parameters for the two contexts as stacked blocks 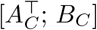, combining the sparse signed interaction matrix *A*_*C*_ with the context-specific baseline drift *B*_*C*_. This highlights that the simulator does not create a new context by simply shifting observed expression values after sampling. Instead, the context enters upstream through the latent dynamics themselves, by altering the regulatory interactions, the baseline drift, or both.

The four UMAP panels in the bottom row then show how these mechanism choices translate into observable geometry. Each point is a simulated cell, color identifies perturbation, and marker shape identifies the context. Under `None` (*A*_1_ = *A*_0_, *B*_1_ = *B*_0_), the two contexts overlap almost completely, as expected when perturbation effects are fully transportable by construction. Under `A`, where only the interaction matrix differs across contexts, the global shape of the manifold is visibly warped, reflecting changes in how perturbation signals propagate through the regulatory network. Under `B`, where only the baseline drift changes, the manifold exhibits a more uniform displacement. Under `Both`, the geometry is the most strongly separated, indicating the largest departure from transportability.

##### Benchmarking CausalDGP with real Perturb-seq data

We contrast CausalDGP with real Perturb-seq data (per cell context) along marginal, co-expression, perturbation-effect, effective diversity, and discriminability metrics (Table 6). Notably, CausalDGP falls within observed ranges for dispersion (CoV), sparsity (Zero), co-expression (Frac|*r*|>0.3), and effect size 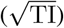.

**Table 6:**
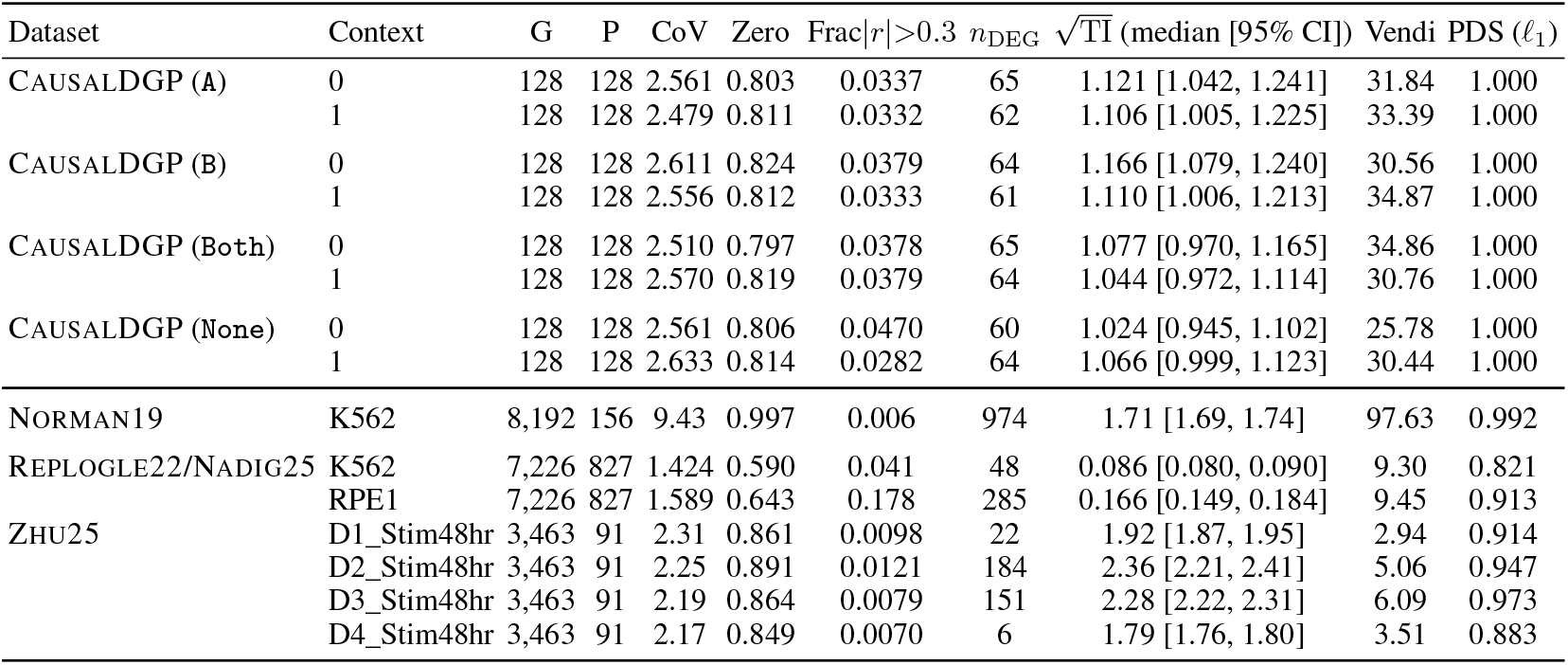
Perturbation effect, effective diversity, discriminability, and marginal statistics per cell-line context. **G**: number of genes; **P**: number of perturbations; **CoV**: gene-wise CoV (dispersion); **Zero**: gene-wise zero proportion (sparsity); **Frac**|*r*|>0.3: fraction of gene pairs with absolute Pearson correlation > 0.3 (metacell-aggregated); *n*_DEG_: DEGs per perturbation; 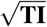 square root of *noise adjusted* transcriptome-wide impact [Nadig et al., 2025], reported as median [95% CI]; **Vendi**: pseudobulk Vendi diversity score; **PDS** (*l*_1_): perturbation discrimination score. Marginal statistics/effect values are bootstrap medians; Vendi and PDS (*l*_1_) (estimated by split-half self-consistency ) are means over 5 seeds.

##### Remarks

Our theoretical analysis assumes a linear SDE (2), whereas real perturbation responses involve nonlinear, multilayered regulatory cascades [Amit et al., 2009]. Linearity is confined to the latent drift; although the emission model uses a gene-wise nonlinear softplus link with a negative binomial observation model, CausalDGP does not explicitly represent nonlinear multi-gene synergies. Context-specific mechanisms are encoded by *A*_*C*_ and *B*_*C*_, while the intervention is represented by Γ_*q*_. This tractable formulation allows the effects of causal mechanism shifts to be precisely characterized, and the simulated cross-context generalization gaps are consistent with those observed in real Perturb-seq data. Context-dependent intervention effects could be incorporated by allowing Γ_*q*_ to depend on *C*, while nonlinear regulatory interactions would require replacing the linear drift with a nonlinear transition function. Existing real-data benchmarks lack ground-truth causal structure, while semi-synthetic simulators such as CausalDGP provide it, albeit under simplified dynamics and observation models. Importantly, the central conclusion remains unchanged: transportability depends on whether the causal mechanisms governing perturbation responses are invariant across contexts.

## C Datasets

### C.1 Perturb-seq dataset pre-processing

Table 7 summarizes the pre-processing pipeline. We follow Dibaeinia et al. [2026] for Zhu25 and Mejia et al. [2025] for Norman19 and Replogle22/Nadig25.

**Table 7:** Dataset-specific pre-processing and filtering criteria for real-world Perturb-seq datasets.

| Criterion | NORMAN19 | REPLOGLE22/NADIG25 | ZHU25 |
| --- | --- | --- | --- |
| *Cell-level QC* |  |  |  |
| Minimum detected genes per cell | 200 | 200 | 100 |
| Total UMI filter | Source-study QC<sup>a</sup> | Source-study QC<sup>a</sup> | $[\max\{1,400, m - 9 \text{ MAD}\}, m + 9 \text{ MAD}]$ |
| *Gene-level QC* |  |  |  |
| Minimum gene abundance | $\geq 3$ cells | $\geq 3$ cells per pair | $\geq 100$ cells and $\geq 100$ total UMIs |
| *Feature selection* |  |  |  |
| Selection scope | All retained cells | Each K562-anchored pair | Each time point; donors pooled |
| HVG | 8,192 | 7,226 (RPE1) / 7,595 (HepG2) / 7,563 (Jurkat) | 3,463 |
| *Perturbation filtering* |  |  |  |
| Minimum cells per perturbation | 256 | 64 | 64 |
| Knockdown-ratio filter | N/A | Source-study QC<sup>a</sup> | $< 0.5$ at guide and cell levels<sup>b</sup> |
| Minimum donors per time point | N/A | N/A | $\geq 2$ at every time point |
| *Dataset summary* |  |  |  |
| Analysis contexts | 1 (K562) | 4 cell lines in 3 K562-anchored pairs | 12 donor-timepoint combinations ($4 \times 3$) |
| Perturbation design | Single- and dual-gene (CRISPRa) | Single-gene (CRISPRi) | Single-gene (CRISPRi) |

<sup>a</sup> No additional filter of this type was applied; source-study QC was retained.
<sup>b</sup> Ratios compare target-gene expression with the matched donor-timepoint control mean.
Final matrices were normalized to $10^4$ counts per cell and $\log_1\text{p}$ -transformed. HVG, highly variable gene; MAD, median absolute deviation; UMI, unique molecular identifier.

### C.2 Context-aware splitting strategy

#### Shared pre-split planner

Before constructing either final split, we run a shared planning step. The planner constructs provisional splits for both strategies, selects the perturbations to hold out in the target context(s), records the within-context test-set size for each context-perturbation group, and computes the overall training-set size under both schemes. These quantities are then fed into the subsequent fairness-alignment stage. This shared planner ensures that the final within-context and cross-context splits are directly comparable, rather than reflecting different random choices or different effective data budgets.

#### Within-context split

For each (perturbation, context) combination (*q, c*), cells are split 50*/*20*/*30% into train, validation, and test sets. The model sees every pair during training but is evaluated on held-out cells.

#### Cross-context split

Given a target context *c*_test_, we randomly hold out 50% of its non-control perturbations denoted *Q*_held_, from training, creating unseen (perturbation, context) combinations (*q, c*_test_) for *q* ∈ *Q*_held_. The remaining perturbations in (*q*^*′*^, *c*_test_) : *q*^*′*^ ∉ *Q*_held_ and all perturbations in other contexts, including _held_ itself, form the training pool. Control cells in *c*_test_ are split 50*/*20*/*30% as in the within-context regime, preserving held-out controls for evaluation.

#### Fairness alignment

To ensure the two regimes are directly comparable, we apply three post-hoc alignment steps (marked “Drop” in Figure 7):

i. *Perturbation alignment*: the within-context test set in *c*_test_ is restricted to the same held-out perturbations *Q*_held_, so that both regimes are evaluated on identical (*q, c*_test_) combinations.
ii. *Test-size alignment*: cross-context test sets are downsampled to match the within-context test-set size per perturbation.
iii. *Train-size alignment*: the larger training set is downsampled to match the smaller.

## D Metrics

### D.1 Effective perturbation response diversity: Vendi score

Table 8 details the null-calibration hyperparameters and how they are determined. Calibration is performed separately within each evaluation context.

**Table 8:** Null-calibration hyperparameters for the pseudobulk Vendi score.

| Symbol | Parameter | Value | Summary |
| --- | --- | --- | --- |
| $n_{\text{null}}$ | Control split size | $\min \left\{ \left\lfloor \text{median}_q(N_q) \right\rfloor, \left\lfloor N_0/2 \right\rfloor \right\}$ | Approximates the typical empirical perturbation sample size. The second term ensures that two disjoint control subsets can be formed. |
| $S$ | Number of null splits | 200 | Number of independently generated pairs of disjoint control subsets. Cells may be reused across different splits. |
| $\rho$ | Null-distance quantile | 0.95 | Uses the 95th percentile of the control split distances as a conservative estimate of finite-sample null variation. |
| $\kappa$ | Target null similarity | 0.95 | Sets the RBF scale so that a distance equal to $\delta_\rho$ has similarity 0.95. |
| $\sigma_{\text{null}}^2$ | Derived RBF variance | $-\frac{\delta_\rho}{2 \log \kappa}$ | Ensures $\exp[-\delta_\rho/(2\sigma_{\text{null}}^2)] = \kappa$. |

### D.2 Perturbation effect metrics

#### Perturbation effect discrimination score (PDS)

To assess whether each predicted perturbation effect is closer to its own ground truth than to those of other perturbations, we adopt the PDS of Wu et al. [2025]. For each perturbation *q, r*_*q*_ measures the fraction of competing predictions that the matched prediction 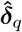 outranks:

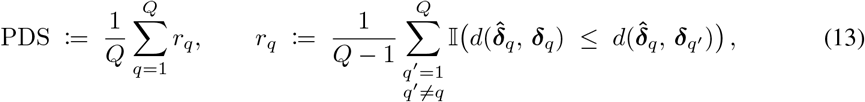

where 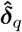 is the predicted perturbation effect (7), ***δ***_*q*_ is the corresponding observed effect, and *d* is a distance function (we report *l*_1_, but *l*_2_ and cosine variants could also be considered). A perfect model places every prediction closest to its own target, yielding PDS = 1; a mode-collapsed model that emits near-identical profiles across perturbations achieves PDS ≈ 0.5, the expected value under random assignment. PDS therefore complements per-perturbation metrics such as Pearson ***δ*** and 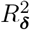, which can remain high when a model captures the shared control-to-perturbation shift but fails to distinguish individual perturbation effects. Unlike the Vendi score, which requires no ground truth, PDS requires matched observations and is applicable only at evaluation time. Together, the Vendi score and PDS provide complementary diagnostics: the former measures whether predictions are *diverse*, the latter whether they are *correctly matched* to their targets.

#### **Pearson *δ* and** 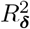

For each perturbation *q*, we compute gene-level effects ***δ***_*q*_ and 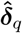 from the observed and predicted mean profiles, respectively. The Pearson correlation between 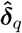 and ***δ***_*q*_ across genes measures whether the model recovers the correct *direction* of perturbation effects irrespective of their magnitude:

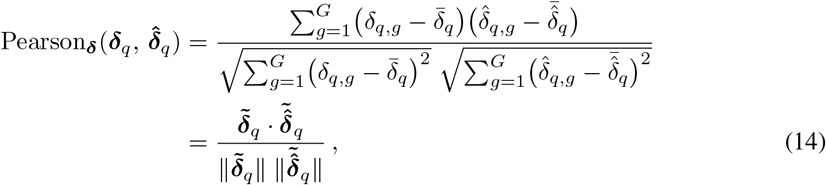

where 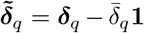. The coefficient of determination 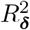 additionally penalizes miscalibrated effect sizes:

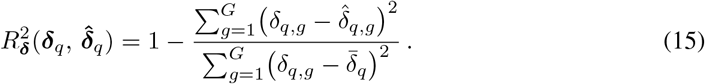

A model that predicts effects proportional to the truth 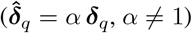 achieves perfect Pearson ***δ*** but 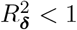. Both metrics are computed across all *G* genes and, separately, restricted to differentially expressed genes (DEGs) identified by a Welch *t*-test with Benjamini–Hochberg correction [Welch, 1947, Benjamini and Hochberg, 1995] at significance level *α*_FDR_ = 0.05. We report the median over perturbations. Because both metrics are evaluated per perturbation and then aggregated, a mode-collapsed model that predicts the population-average effect can still attain moderate Pearson ***δ*** whenever perturbation effects share a common direction, though 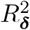 is more sensitive to this failure. This motivates the complementary use of PDS (13) and VR (8).

#### Transcriptome-wide impact (TI)

Following the transcriptome-wide impact (TI) framework proposed by Nadig et al. [2025], we combine `DESeq2` [Love et al., 2014] with adaptive shrinkage (`ash`) [Stephens, 2017] to estimate the noise-adjusted distribution of perturbation effects. Counts are summed within each condition and replicate to form pseudobulks. For gene *g* and pseudobulk *i*, DESeq2 fits

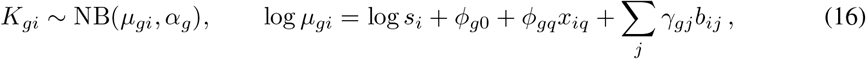

where *s*_*i*_ is size factor estimated by median-of-ratios by default, *x*_*iq*_ indicates perturbation *q, b*_*ij*_ indicates batch *j*, and *α*_*g*_ is the gene-specific dispersion. The perturbation-versus-control contrast yields a log_2_ fold-change estimate 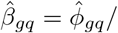 ln 2 and standard error 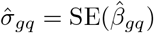, with sampling model

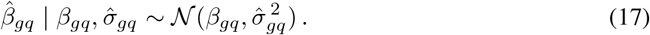

To separate biological effects from estimation noise, `ash` models the centered true effects across genes using a zero-centered normal mixture,

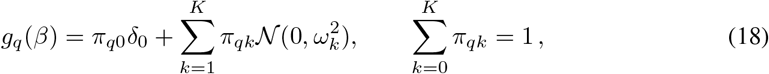

and estimates the mixture weights by marginal maximum likelihood. TI is the variance of the fitted effect-size distribution,

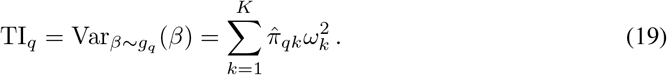

Thus, TI is measured in squared log_2_-fold-change units, whereas 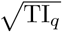 is in log_2_-fold-change units.

### D.3 DEG-level set metrics

Beyond weighting gene-level scores by DEG membership, we treat differential expression as a binary classification problem. For each perturbation *q*, let 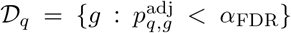 denote the set of observed DEGs, where 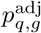 is the Benjamini–Hochberg-adjusted *p*-value from a Welch *t*-test comparing single-cell expression of gene *g* between *P (Y*_*g*_ |do(*Q*=*q*), *C)* and *P (Y*_*g*_ | do(*Q*=0), *C)*, with *α*_FDR_ = 0.05. Let 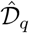 denote the analogous set obtained by applying the same test to predicted single-cell profiles. Standard set-overlap metrics then quantify whether the model preserves the combinatorial identity of each perturbation:

- *Jaccard similarity:* 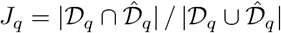,
- *Precision:* 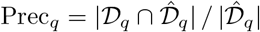,
- *Recall:* 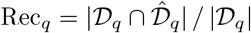 .

Recall, also called *top-k DEG recall* when 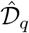 is defined by the *k* most significant predicted genes, is the variant adopted by Wu et al. [2025]. Because DEG-set metrics require predicted single-cell profiles to estimate cell-to-cell variance, we compute them only for single-cell predictors. These metrics complement the continuous scores above: a model can achieve high Pearson ***δ*** by capturing large effects while missing weaker but biologically significant genes, a failure that low recall exposes. We report the median of each metric over perturbations.

### D.4 Reconstruction metrics

#### Mean absolute and mean squared error

We report point-wise reconstruction error on raw counts. Let 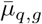 denote the empirical mean expression of gene *g* among cells receiving perturbation *q*, and let 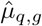 denote the corresponding model prediction. For perturbation *q*:

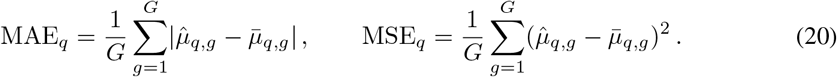

Unlike Pearson ***δ*** and 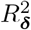, these errors are computed on absolute expression levels rather than shifts from control and are hence sensitive to baseline calibration, *i*.*e*., unperturbed gene expression contributes to the error independently of perturbation-effect accuracy. As with the effect-based scores, both are evaluated across all *G* genes and restricted to DEGs, and we report the median MAE over perturbations.

#### Distributional distances

The pseudobulk metrics above collapse each perturbation population to a single mean vector, discarding cell-to-cell heterogeneity. Distribution-level metrics retain this information and fall into two complementary classes. Single-cell distributional distances (MMD, FD, and JSD) are applicable only to predictors that produce single-cell outputs.

##### Parametric (raw counts)

Because scRNA-seq counts are sparse, overdispersed, and often heavy-tailed, we fit separate negative-binomial distributions to the observed and predicted count vectors for each gene and quantify their discrepancy using the Jensen–Shannon divergence (JSD) [Lin, 2002]. Because the JSD between two negative-binomial distributions has no closed-form expression, we approximate it by summing their probability mass functions over a truncated discrete support, with the upper bound determined from the ( 1 *−* 10^−8^) quantiles of the fitted distributions and the maximum empirical count. The truncated probability masses are then renormalized before computing the JSD. This deterministic procedure avoids Monte Carlo error and respects the discrete, overdispersed nature of the data, but depends on the assumed distributional family and remains sensitive to library-size normalization.

##### Non-parametric (latent embedding)

An alternative projects both observed and predicted cells into a lower-dimensional representation, such as PCA or a foundation-model embedding (*e*.*g*., scGPT [Cui et al., 2024]), and compares the resulting point clouds using a distributional distance. Common choices include the maximum mean discrepancy, 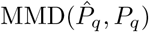, with an RBF kernel [Gretton et al., 2012]; the *p*-Wasserstein distance [Villani, 2009]; and the Fréchet distance (FD), computed from the first two moments of the embedding [Heusel et al., 2017]. These metrics are agnostic to the count distribution but depend on the quality and dimensionality of the embedding. As with the Vendi score (Algorithm 1), we compute the non-parametric distances in PCA space. For both MMD and FD, we fit a single shared PCA projection to the pooled observed control cells within each evaluation context. We then use this fixed projection to transform the observed and predicted cells for every perturbation condition. The distance is computed separately for each perturbation in this common PCA space and subsequently aggregated across perturbations.

## E Full experimental results

### E.1 Reproducing the experiments

All experiments were conducted on a SLURM-managed high-performance computing cluster using GPU-enabled compute nodes. Data generation, loading, and pre-processing used multi-process CPU workers (Intel Xeon Gold 6448Y), while model training and inference used GPU acceleration (NVIDIA L40S, 48 GB). Each job was allocated sufficient system memory (64 GB for synthetic datasets and 256 GB for real datasets) to support parallel workers and GPU execution. Runtimes generally ranged from several hours to approximately 3 days for the largest configurations over 10 runs.

The linearPCA baseline is a bilinear model of the form 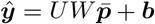, where *U* contains the PCA gene embeddings, 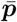 is the mean embedding of the perturbation’s target genes, and *W* is estimated using two-sided ridge regularization with *λ* = 0.1 (Table 4). The gene embeddings are obtained by applying full-SVD PCA with up to 10 components to the transpose of the matrix of training perturbation-level pseudobulks, treating genes as observations and perturbations as features.

Table 9 summarizes the architectural and training hyperparameters used by the deep learning models.

**Table 9:** Architecture and training hyperparameters for deep learning models.

| Model | Architecture | Training |
| --- | --- | --- |
| scVI | Three-layer, width-128 MLP encoder and decoder; latent dimension 10; ReLU and batch normalization; encoder dropout 0.1; ZINB likelihood with gene-wise dispersion. | Adam (lr = $10^{-3}$, wd = $10^{-6}$); batch 512; up to 800 epochs; KL warmup 400 epochs; early-stopping patience 45. |
| GEARS | One co-expression and one perturbation-graph SGConv layer, using GO when available and co-expression otherwise; hidden width 64; shared recovery MLP $64 \rightarrow 128 \rightarrow 64$ for residual gene-space prediction; ReLU and batch normalization; uncertainty head predicting gene-wise log variance. | Adam (lr = $10^{-3}$, wd = $5 \times 10^{-4}$); batch 64; 20 epochs; StepLR ($\gamma = 0.5$ per epoch); gradient-value clipping at 1. |
| CPA | Variational latent dimension 128; hidden width 256; encoder/decoder configured with three layers; perturbation and context embeddings configured with two and one layers, respectively; ReLU; dropout 0.1; Gaussian/MSE reconstruction. | Two Adam optimizers (lr = $10^{-3}$, wd = $10^{-8}$); batch 128; up to 100 epochs; early-stopping patience 10; six warmup epochs; generator-to-adversary update ratio 1:6. |
| STATE | Two-layer basal and perturbation MLPs with width 64; width-64 bidirectional GPT-2 with two blocks, four heads, and FFN width 256; GELU; dropout 0.1; residual gene-space prediction. | Adam (lr = $3 \times 10^{-4}$, wd = 0); batch of four sets of $S$ cells; up to 100 epochs; early-stopping patience 10; energy-distance loss plus 0.1 context-classification and $10^{-5}$ residual L1 penalties. |
| scLDM VAE | Eight encoder and eight decoder blocks; width 32; eight self-attention and four cross-attention heads; 16 latent tokens of width 16; SwiGLU; dropout 0; NB decoder. | AdamWLegacy (lr = $10^{-3}$, $\beta = (0.9, 0.95)$, wd = 0); batch 128; $100 \lfloor N_{\text{cells}}/128 \rfloor$ steps; 10% warmup and square-root decay over the final 10%. |
| scLDM DiT | Eight DiT blocks; width 256; eight heads; FFN width 684; 16 latent tokens; adaptive LayerNorm; dropout 0; joint conditioning with CFG dropout 0.8; linear flow matching with velocity prediction. | AdamW (lr = $5 \times 10^{-4}$, wd = 0); same step budget as the VAE; 10% warmup followed by cosine decay; EMA decay 0.9999; adaptive Dopri5 sampling at 50 requested output times, with guidance weight $\omega \in \{1, 5, 10\}$ |

$S$ is the minimum number of cells per perturbation–context group across the training and validation splits. $N_{\text{cells}}$ is the number of scLDM training cells; the stated step budget is applied separately to the VAE and DiT stages.

### E.2 Validation and robustness of the Vendi score

Across 100 randomly simulated DirectDGP datasets (Figure 8) both Vendi and covariance effective rank increased with the number of perturbations, with similar OLS slopes (*β*_1_ = 0.870 and 0.891, respectively). Effective rank showed a substantially stronger Pearson correlation with *P* (*r* = 0.984) than Vendi (*r* = 0.660) because it remained tightly coupled to its maximum value of *P −* 1. As expected, Vendi exhibited downward deviations when perturbation effects were weak or sparse. Thus, the higher correlation for effective rank likely reflects a near-mechanical dependence on the number of perturbation groups, whereas Vendi is more sensitive to whether those perturbations produce distinguishable responses. Moreover, in Gaussian-noise experiments on Norman19 (Figure 9), Vendi and PDS (*l*_1_) remain stable under low-to-moderate noise and decline under severe noise, whereas effective rank increases with noise.

Figure 10 evaluates the sensitivity of the Vendi score to perturbation sample size and finite-sample noise. Under the population null (*ϵ* = 1), severe sample-size heterogeneity increased mean Vendi relative to balanced sampling. This inflation was attenuated at *ϵ* = 1.5 and negligible at *ϵ* ≥ 2. The large-pool mean oracle also yielded a lower Vendi Ratio than the sampling-matched oracle at low effect factors, but this difference diminished with increasing effect factor and sample size, becoming negligible by *N*_*q*_ = 256 at *ϵ* = 2 and at all evaluated sample sizes for *ϵ* = 5. Thus, fixed-size null calibration and deterministic prediction can introduce bias, but their practical impact is concentrated under null or low-effect conditions and is small for moderate-to-strong effects or larger sample sizes.

### E.3 Perturbation response geometry: diversity and discrimination

Figure 11 provides the cross-context counterpart to Figure 3b for the two multi-context datasets, Zhu25 and Replogle22/Nadig25. Relative to within-context evaluation, the dominant change is a leftward shift in PDS, indicating weaker discrimination when models predict unseen perturbation– context combinations. On Zhu25 (circles), all models approach chance-level discrimination (PDS *≈* 0.5) and substantially underestimate response diversity (VR *<* 1). Thus, cross-context predictions on Zhu25 fail to preserve both perturbation identity and observed diversity.

On Replogle22/Nadig25 (triangles), the degradation is milder and model performance separates more clearly. scLDM retains the strongest discrimination, whereas scVI best preserves diversity, with VR close to one. STATE and STATE-Geneformer retain moderate PDS but underestimate diversity, while GEARS and linearPCA remain near chance and exhibit low VR. These results reveal two distinct, model-dependent failure modes under cross-context transfer: loss of perturbation discrimination and loss of response diversity. Strong within-context performance therefore does not necessarily imply robust generalization across biological contexts.

### E.4 Cross-context generalization gap

Appendix Figures 12–15 provide the full per-context results underlying Section 4.3.3 across all three benchmarks. We report five complementary metrics: 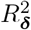 (DEGs-restricted; plotted as 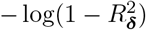 to accommodate its wide range) and DES (recall), where larger values indicate better perturbation-specific agreement; MAE (DEGs-restricted) and MMD, where smaller values are better; and VR, which measures relative effective diversity.

#### E.4.1 Cross-context generalization gap for Zhu25

Figure 12 shows a broad cross-context generalization gap on Zhu25 across all six held-out donor– activation combinations. Under within-context evaluation, STATE, STATE-Geneformer, and scLDM generally perform best across DEG-restricted *R*^2^, DES recall, MAE, and MMD, while CPA remains competitive, particularly in *R*^2^. Under cross-context evaluation, these advantages contract sharply. Across all six contexts, DEG-restricted *R*^2^ decreases and MAE and MMD increase for all four models; DES recall also declines consistently for STATE, STATE-Geneformer, and scLDM. Median *R*^2^ falls below zero, indicating poor perturbation-specific agreement for unseen perturbation–context combinations.

Because these declines recur across all six contexts, they are not driven by a single donor or activation state. Relative performance nevertheless remains metric-dependent: CPA, STATE, and STATE-Geneformer retain the strongest *R*^2^, whereas scLDM retains the highest DES recall and lowest MMD. Weaker methods exhibit less consistent changes on some metrics, partly because their within-context performance is already low.

VR requires a more nuanced interpretation. It changes little for CPA and only moderately for STATE and STATE-Geneformer, but declines substantially for scLDM. Moreover, cross-context VR remains well below one for every model, indicating systematic underestimation of response diversity. Thus, cross-context failure is not explained solely by additional diversity collapse: models also lose alignment between predicted responses and the correct perturbations in held-out biological contexts.

#### E.4.2 Cross-context generalization gap for Replogle22/Nadig25

Figure 13 shows systematic cross-context generalization gaps for RPE1–K562. Although RPE1–K562 is the least concordant Replogle22/Nadig25 pair, its TI correlation remains substantially higher than the cross-donor correlations in Zhu25 (Table 2). Cross-context evaluation therefore reduces performance without eliminating perturbation-specific signal. Performance is metric-dependent. Within context, STATE-Geneformer achieves the strongest DEG-restricted Pearson and 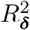, whereas scLDM achieves the strongest PDS (*l*_1_) and lowest MMD; STATE and CPA remain competitive. The context-aware linear baselines are also informative: Context-Average and Context-linearPCA retain moderate DEG-restricted Pearson under transfer, with Context-linearPCA approaching CPA. However, their near-random PDS and negative 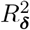 indicate limited perturbation-specific discrimination. Thus, simple context-conditioned summaries capture a transportable component of the average response structure, but not the full perturbation-specific geometry.

Across RPE1–K562, HepG2–K562, and Jurkat–K562, CPA, STATE, STATE-Geneformer, and scLDM exhibit positive transfer gaps in DEG-restricted Pearson and PDS. RPE1–K562, which has the lowest TI concordance (Table 2), consistently exhibits the largest gaps in both metrics (Figure 14). Other metrics show less consistent relationships with TI concordance, with changes in 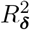, MAE, DES recall, MMD, and VR depending on both model and context pair. Thus, mechanistic similarity aligns most clearly with transfer in DEG-restricted Pearson and PDS.

#### E.4.3 Cross-context generalization gap for CausalDGP

Figure 15 provides a controlled view of cross-context generalization because the source of context shift is known and tunable by construction. The results support the transportability condition in Section 2.3. When neither the baseline state nor the interaction mechanism changes (`None`), CPA, GEARS, and scLDM retain strong cross-context performance, with only small reductions in DEG-restricted Pearson, PDS, and DES recall. Changing only the baseline drift (`B`) produces moderate degradation for CPA and GEARS, whereas scLDM remains comparatively stable.

**Figure 15:**
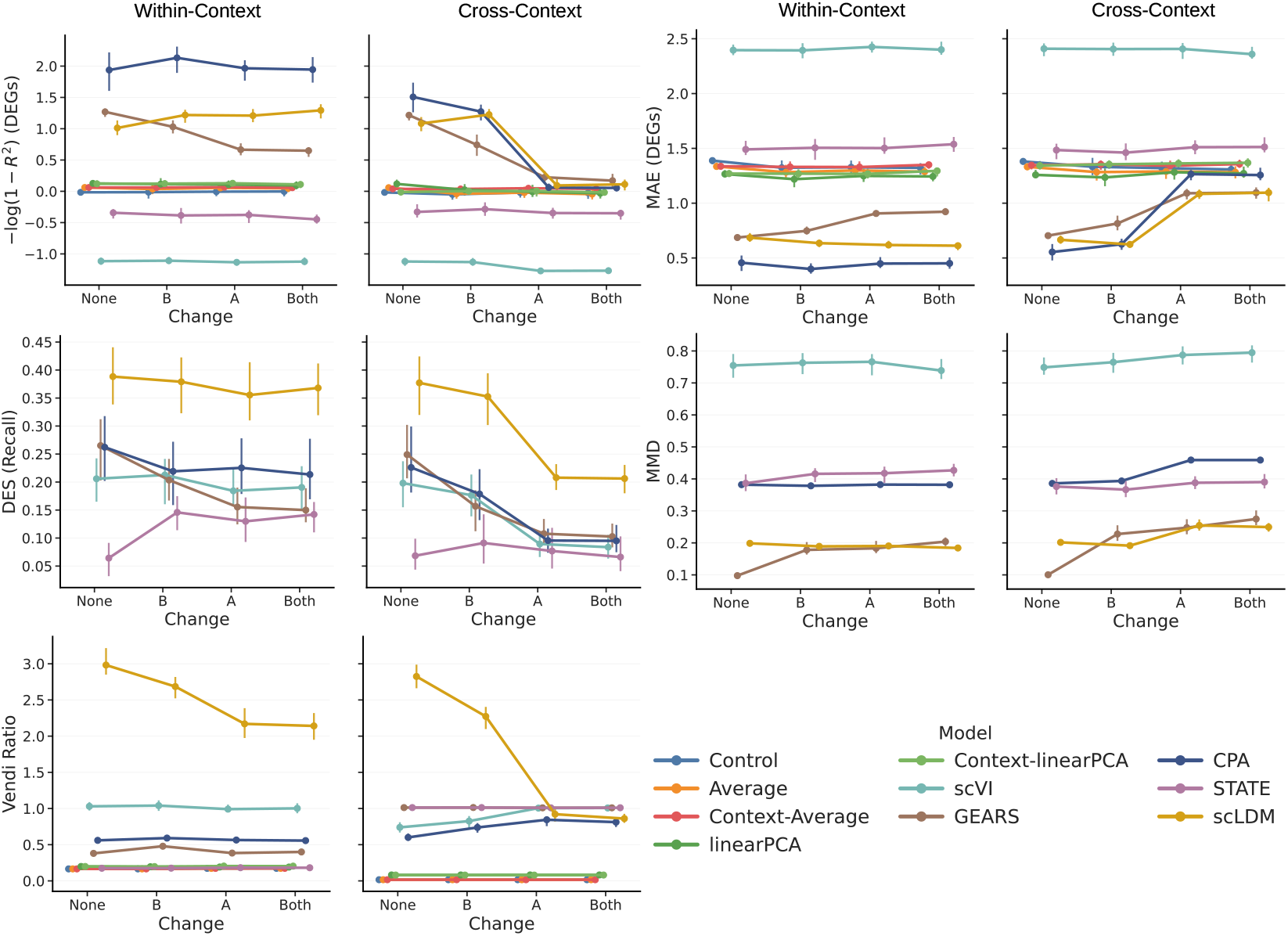
Additional results on the cross-context generalization gap in CausalDGP under within-context and cross-context evaluation. The *x*-axis denotes the four cross-context mechanism modes (`None`, `A`, `B`, `Both`). We plot medians and interquartile ranges. scLDM is evaluated with *ω* = 5.

The pattern changes sharply when the interaction matrix *A*_*C*_ differs across contexts. Under `A` and `Both`, CPA, GEARS, and scLDM exhibit substantially larger transfer gaps: DEG-restricted *R*^2^ and Pearson decline, MAE increases, and PDS and DES recall deteriorate. scLDM nevertheless retains the strongest PDS and DES recall under these shifts. STATE remains near chance in PDS across conditions, limiting its ability to reveal the same ordering because of a floor effect.

For scLDM, VR moves closer to one under `A` and `Both` even as perturbation-specific accuracy deteriorates. Thus, improved diversity calibration does not imply correct perturbation-response alignment, reinforcing that diversity and discrimination capture distinct failure modes. Overall, transfer gaps are smallest under `None`, larger under `B`, and largest under `A` and `Both` for the models that perform well in context. This supports the prediction that changes to the perturbation-response mechanism are more damaging than baseline-state shifts alone.

#### E.4.4 Alignment between theory and experiments

Table 2 grounds the divergence parameter in an empirically measurable quantity: the cross-context concordance of transcriptome-wide impact (TI) induced by the CausalDGP influence matrix, *A*, recapitulates the range observed across real biological contexts. For example, as the divergence parameter increases, CausalDGP reproduces the low cross-donor TI concordance observed in Zhu25 (Figure 4). This low concordance coincides with poor generalization across all models, supporting TI concordance as a real-data proxy for mechanistic divergence. A similar pattern emerges across cell lines: Jurkat and HepG2 exhibit comparably high TI concordance with K562, whereas RPE1 exhibits substantially lower concordance. Correspondingly, RPE1 *→*K562 produces the largest transfer gaps across all four models for both DEG-restricted Pearson ***δ*** and PDS (*l*_1_) (Figure 14). Together, these results support the theoretical prediction in (6) that transfer gaps widen as TI concordance decreases.

A perturbation-level analysis of STATE on RPE1 →K562 further localizes this aggregate transfer penalty. Table 10 shows that the cross-context penalty is concentrated among a small, reproducible subset of perturbations. Moreover, perturbations with more divergent effects exhibit larger cross-context generalization gaps, reinforcing the relationship between effect divergence and the reduced TI concordance reported in Table 2.

**Table 10:** STATE on Replogle22/Nadig25 (RPE1 *→*K562), for the 532 perturbations held out in *≥* 5 of 10 seeds. We test whether each perturbation’s mean transfer gap (PDS (*l*_1_)) differs from zero (variance-stabilized and BH FDR-controlled). No perturbation shows a corroborated *positive* gap.

| Class | $n$ | % of perts | Median gap | Mean gap | $\sum$ gap (% of total) |
| --- | --- | --- | --- | --- | --- |
| Consistent negative (transfer failure) | 57 | 10.7% | -0.153 | -0.239 | -13.61 (55%) |
| No significant gap | 475 | 89.3% | -0.011 | -0.024 | -11.18 (45%) |
| All analyzed | 532 | 100% | -0.016 | -0.047 | -24.79 (100%) |

### E.5 scLDM classifier-free guidance sensitivity analysis

Figures 16–18 compare classifier-free guidance weight *ω* ∈ {1, 5, 10}, revealing metric-dependent effects. On CausalDGP, stronger guidance improves cross-context PDS (*l*_1_) when *A* changes, but reduces cross-context Pearson ***δ***. For RPE1–K562, PDS improves while cross-context Pearson ***δ*** changes little. On Zhu25, guidance improves in-context Pearson ***δ*** and PDS, but their cross-context values remain near zero and chance, respectively. Across datasets, stronger guidance increases both VR and MMD, indicating that greater response diversity does not necessarily improve distributional fidelity. Thus, tuning *ω* changes the trade-offs between metrics but does not eliminate the cross-context generalization gaps.

**Figure 16:**
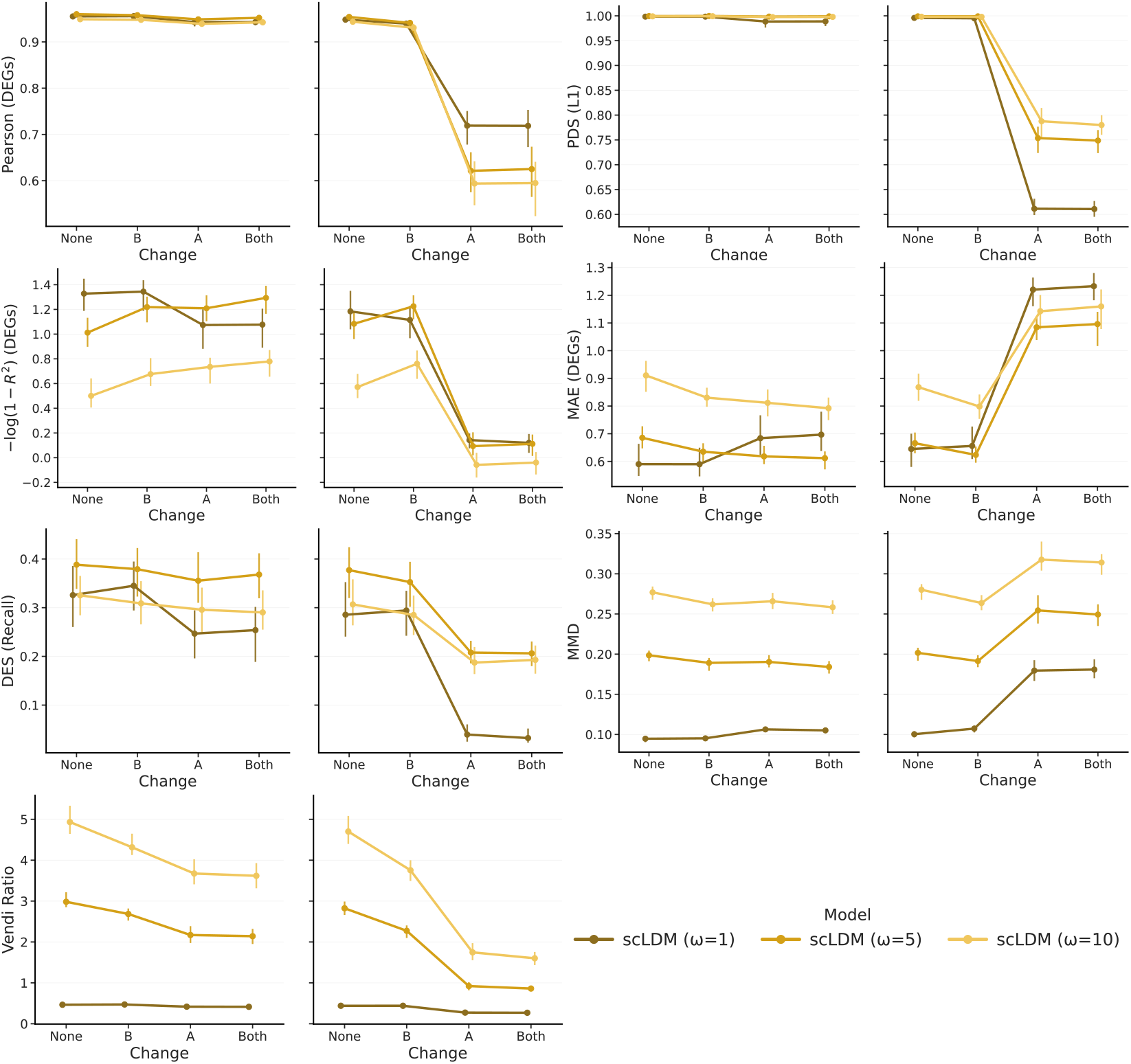
Sensitivity of scLDM to the classifier-free guidance weight *ω* on CausalDGP.

**Figure 17:**
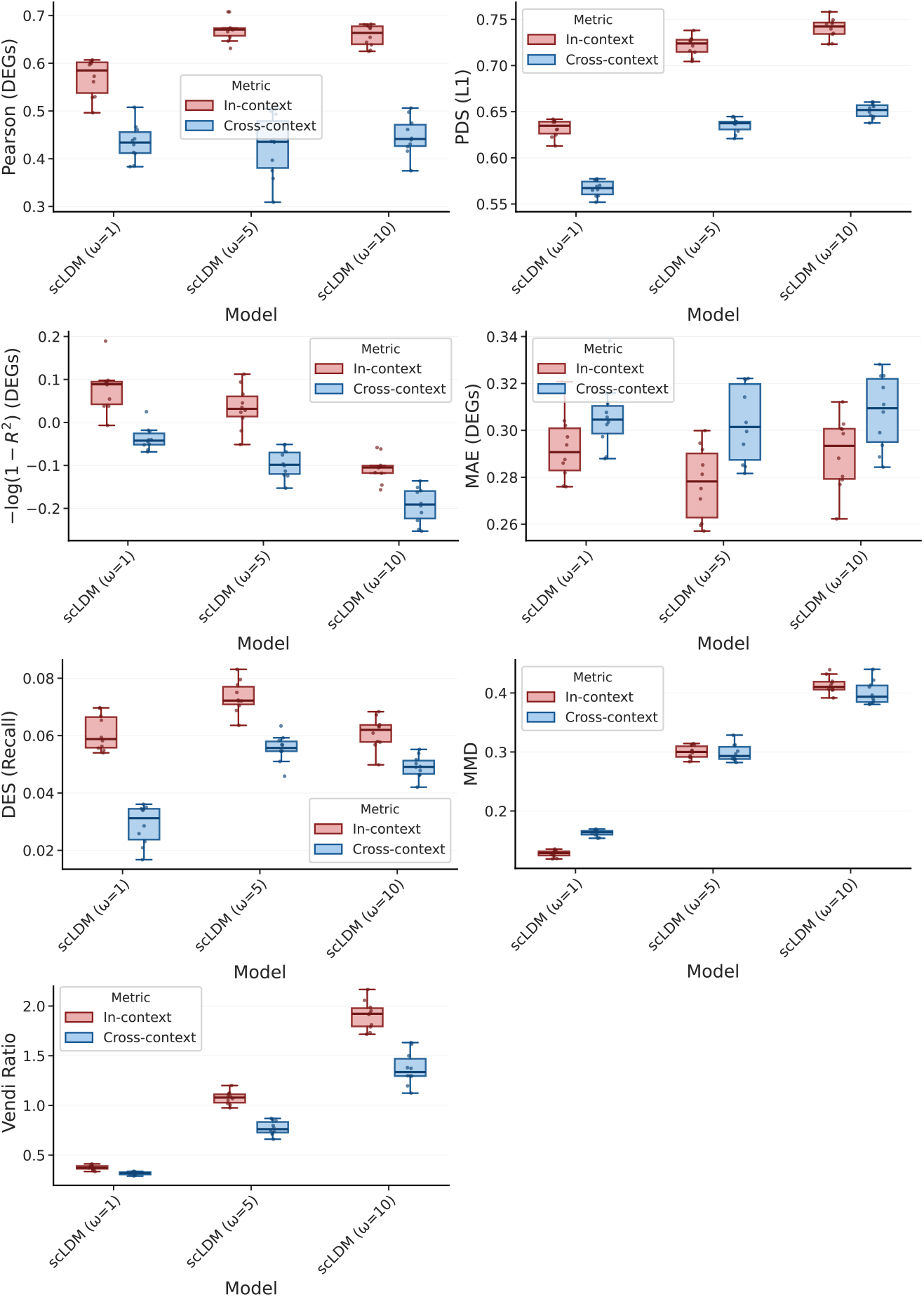
Sensitivity of scLDM to the classifier-free guidance weight *ω* onReplogle22/Nadig25.

**Figure 18:**
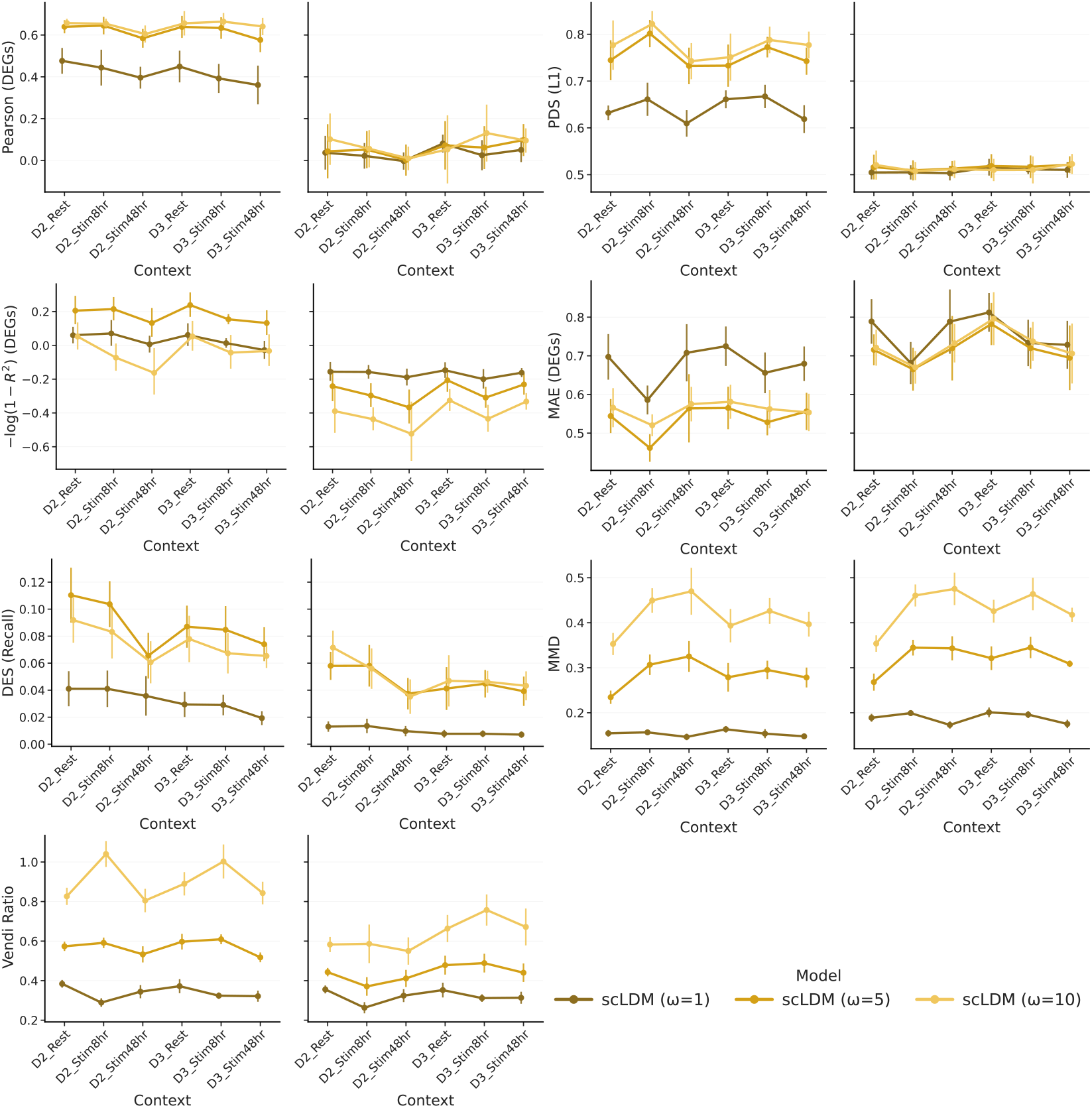
Sensitivity of scLDM to the classifier-free guidance weight *ω* on Zhu25.

Table 11 also summarizes the sensitivity of scLDM to *ω*. Overall, PDS (*l*_1_) increases with guidance strength, whereas Pearson ***δ*** (DEG) generally improves from *ω* = 1 to *ω* = 5 before plateauing or declining at *ω* = 10. These results reveal a trade-off between perturbation discriminability and per-gene fidelity under stronger guidance. Stronger guidance increases predicted effective diversity. Guidance weight *ω* = 5 offers a useful overall compromise, whereas *ω* = 10 often overshoots empirical diversity, although the best-calibrated setting varies across context pairs.

**Table 11:** Sensitivity of scLDM to the classifier-free guidance weight *ω* across three K562-anchored Replogle22/Nadig25 cell-line pairs. Values are reported as median [Q1, Q3] over 10 trials for PDS (*l*_1_), Pearson ***δ*** (DEG-restricted), Vendi ratio (VR). Cross-context performance measures generalization to K562 perturbations withheld from K562 training but observed in the paired source cell line.

| Source | Model | PDS ($\ell_1$) | | Pearson $\delta$ (DEG) | | VR | |
| --- | --- | --- | --- | --- | --- | --- | --- |
|  |  | In-context | Cross-context | In-context | Cross-context | In-context | Cross-context |
| RPE1 | scLDM ($\omega = 1$) | 0.635 [0.626, 0.639] | 0.567 [0.560, 0.574] | 0.585 [0.538, 0.602] | 0.434 [0.412, 0.456] | 0.372 [0.363, 0.390] | 0.318 [0.305, 0.328] |
| | scLDM ($\omega = 5$) | 0.724 [0.715, 0.728] | 0.638 [0.631, 0.639] | 0.671 [0.658, 0.674] | 0.435 [0.381, 0.479] | 1.079 [1.027, 1.112] | 0.761 [0.727, 0.833] |
| | scLDM ($\omega = 10$) | 0.742 [0.734, 0.747] | 0.652 [0.645, 0.657] | 0.664 [0.640, 0.677] | 0.441 [0.427, 0.471] | 1.922 [1.795, 1.979] | 1.336 [1.295, 1.470] |
| HepG2 | scLDM ($\omega = 1$) | 0.693 [0.684, 0.695] | 0.629 [0.623, 0.631] | 0.685 [0.667, 0.705] | 0.584 [0.531, 0.611] | 0.263 [0.250, 0.267] | 0.217 [0.205, 0.228] |
| | scLDM ($\omega = 5$) | 0.787 [0.782, 0.797] | 0.727 [0.720, 0.736] | 0.758 [0.748, 0.763] | 0.666 [0.631, 0.700] | 0.870 [0.784, 0.890] | 0.614 [0.589, 0.670] |
| | scLDM ($\omega = 10$) | 0.804 [0.794, 0.812] | 0.739 [0.735, 0.745] | 0.734 [0.727, 0.744] | 0.677 [0.639, 0.694] | 1.394 [1.278, 1.436] | 0.994 [0.959, 1.105] |
| Jurkat | scLDM ($\omega = 1$) | 0.654 [0.648, 0.656] | 0.604 [0.598, 0.613] | 0.630 [0.620, 0.649] | 0.483 [0.465, 0.519] | 0.405 [0.390, 0.416] | 0.355 [0.341, 0.367] |
| | scLDM ($\omega = 5$) | 0.751 [0.749, 0.756] | 0.697 [0.690, 0.702] | 0.729 [0.719, 0.744] | 0.645 [0.591, 0.660] | 1.227 [1.203, 1.272] | 1.056 [0.846, 1.091] |
| | scLDM ($\omega = 10$) | 0.766 [0.762, 0.773] | 0.711 [0.704, 0.715] | 0.704 [0.700, 0.720] | 0.645 [0.609, 0.662] | 2.221 [2.106, 2.332] | 1.899 [1.462, 1.961] |

We therefore use *ω* = 5 as a common guidance setting in all other experiments.

### E.6 Metric distortion

Figure 19 evaluates *metric distortion, i*.*e*., whether an evaluation metric varies systematically with a nuisance property of the data rather than purely reflecting model quality. In panel (a), the nuisance factor is the control bias (*β*), while in panels (b) and (c), it is the observed DEG percentage. If a metric changes substantially as these nuisance factors vary, part of its value may reflect dataset characteristics such as bias strength or perturbation magnitude rather than genuine predictive performance.

**Figure 19:**
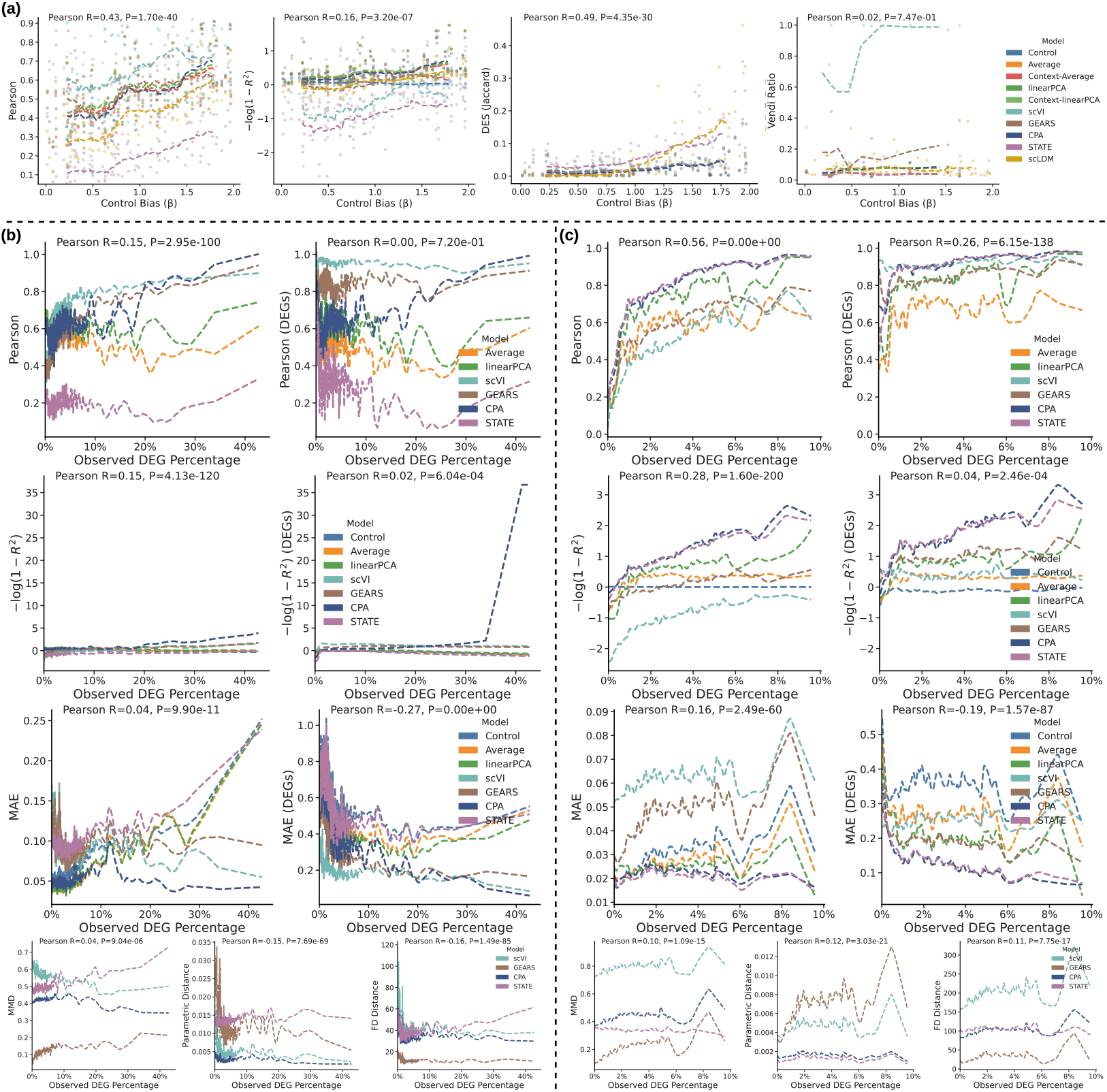
Metric distortion on DirectDGP and Norman19. Points show individual runs, coloured by model; dashed curves show per-model LOESS-smoothed trends. Top annotations report the Pearson correlation (*R*) and two-sided *p*-value (*P* ) computed from all raw points in each panel. (a) On DirectDGP: Pearson ***δ***, 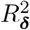, DES (Jaccard), and Vendi Ratio (VR) versus control bias (*β*) across 100 random parameter draws. (b) On DirectDGP: Pearson ***δ***, 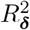, and MAE on all genes and on DEGs, together with MMD, parametric distance, and FD, versus observed DEG percentage across all perturbations and 100 random parameter draws. (c) Corresponding results on Norman19, aggregated across all perturbations over 10 repeated experiments.

In panel (a) each point represents one evaluation result for a given model under one sampled setting, and the dashed curve shows a model-specific locally estimated scatterplot smoothing (LOESS) trend that summarizes how the metric changes with the nuisance factor. In the remaining panels, we omit individual points to avoid visual clutter and show only the smoothed trends.

The annotation at the top of each subplot reports the Pearson correlation (*R*) between the metric and the *x*-axis variable, computed over all raw points in that panel, together with its two-sided *p*-value. A robust metric should exhibit relatively flat LOESS curves and a small absolute correlation, indicating that it is not tracking the nuisance variable. Because these panels contain many observations, very small *p*-values can arise even for weak effects. We therefore place greater emphasis on the magnitude of the correlation and the visual slope of the smoothed curves than on statistical significance alone.

#### Metric distortion with respect to control bias on DirectDGP

In DirectDGP (Algorithm 2), control bias is introduced by the additive term *βλ*_*g*_ in the perturbed mean 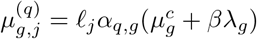. Here *λ*_*g*_ is a gene-specific bias pattern estimated from the real dataset, and *β* controls its strength. Because this term is shared across all perturbations, it induces a common control-referenced shift that is not specific to any individual perturbation. Consequently, larger *β* can make simulated data appear easier to predict even without actually improving recovery of perturbation-specific effects. We therefore prefer metrics that are insensitive to control bias; otherwise, a metric may reward nuisance structure in the simulator rather than genuine model quality, leading to unfair comparisons.

In panel (a) of Figure 19, Pearson ***δ***, 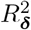, and DES (Jaccard) all show noticeable dependence on control bias *β*, with correlations of *R* = 0.43, 0.16, and 0.49, respectively (all *P ≤* 1 *×* 10^−6^). Their LOESS curves also drift visibly upward as *β* increases, even for simple baselines such as Control or Average. This indicates that these metrics improve systematically as control bias strengthens, meaning their values are partly driven by a nuisance property of the data-generation process. By contrast, VR is the only metric in panel (a) that remains largely insensitive to control bias, with near-zero and non-significant correlation (*R* = 0.02, *P* = 0.747) and comparatively flat trends across models. This suggests that VR is substantially more robust to control bias than the other metrics considered.

#### Metric distortion with respect to observed DEG percentage

Panels (b) and (c) of Figure 19 examine distortion with respect to DEG percentage on DirectDGP and Norman19, respectively. A strong correlation with the observed DEG percentage is undesirable because it implies that the metric is partly measuring perturbation strength or task difficulty rather than prediction quality alone. Such dependence can confound comparisons across perturbations, since models may appear better simply because the underlying perturbation affects a larger fraction of genes. We therefore prefer metrics that are less sensitive to DEG percentage.

For both DirectDGP and Norman19, the DEG-restricted versions of Pearson ***δ*** and 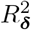 are consistently more robust than their all-gene counterparts. On DirectDGP, Pearson ***δ*** computed on all genes has a positive correlation with observed DEG percentage (*R* = 0.15), whereas the DEG-only version is essentially uncorrelated (*R* = 0.00). A similar pattern holds for 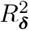: the all-gene version has *R* = 0.15, while the DEG-only version is much less sensitive (*R* = 0.02). On Norman19 the same trend is even clearer. Pearson ***δ*** on all genes shows strong dependence on DEG percentage (*R* = 0.56), whereas the DEG-only version reduces this substantially (*R* = 0.26). Likewise, 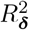 drops from *R* = 0.28 (all genes) to *R* = 0.04 (DEG-restricted). These results indicate that restricting evaluation to DEGs reduces metric distortion and yields a fairer assessment across perturbations with different response magnitudes.

In contrast, MAE is not robust in either formulation. On DirectDGP, both all-gene and DEG-only MAE vary systematically with DEG percentage, and on Norman19 both versions also remain sensitive to this nuisance factor. Unlike Pearson ***δ*** and 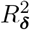, switching from all genes to DEGs does not resolve the distortion problem for MAE. This suggests that MAE remains entangled with perturbation magnitude and is therefore less suitable when robust evaluation across heterogeneous perturbations is desired.

Finally, the distance-based metrics in the bottom row of panels (b) and (c) are generally less distorted than several of the gene-wise metrics. Among them, MMD appears the most robust, showing the weakest dependence on observed DEG percentage. Parametric distance and FD exhibit somewhat larger drift with DEG percentage in at least some settings. Taken together, these results suggest that MMD is the most insensitive of the distance-based metrics to variation in perturbation strength.

## F Broader impacts

This work provides a framework for identifying when perturbation predictions are unreliable across biological contexts, which can help prevent misallocation of experimental resources in drug development. We caution that inflated within-context metrics may mask cross-context failures. Hence model predictions should not guide therapeutic decisions without independent validation. We do not foresee direct negative societal consequences from this evaluation framework.

## G Declaration of LLM usage

We used large language models (LLMs) to assist with code used to prepare plots and tables and to improve the readability of author-written manuscript text and checklist responses. All LLM-assisted code and text were reviewed by the authors. Code was tested and its outputs were inspected, while language edits were checked to preserve the intended scientific meaning. We take responsibility for the final content of this work, including all text, claims, and artifacts produced with the aid of LLMs.

